# Cdc13 exhibits dynamic DNA strand exchange in the presence of telomeric DNA

**DOI:** 10.1101/2023.12.04.569902

**Authors:** David G. Nickens, Zhitong Feng, Jiangchuan Shen, Spencer J. Gray, Robert H. Simmons, Hengyao Niu, Matthew L. Bochman

## Abstract

Telomerase is the enzyme that lengthens telomeres and is tightly regulated by a variety of means to maintain genome integrity. Several DNA helicases function at telomeres, and we previously found that the *Saccharomyces cerevisiae* helicases Hrq1 and Pif1 directly regulate telomerase. To extend these findings, we are investigating the interplay between helicases, single-stranded DNA (ssDNA) binding proteins (ssBPs), and telomerase. The yeast ssBPs Cdc13 and RPA differentially affect Hrq1 and Pif1 helicase activity, and experiments to measure helicase disruption of Cdc13/ssDNA complexes instead revealed that Cdc13 can exchange between substrates. Although other ssBPs display dynamic binding, this was unexpected with Cdc13 due to the reported *in vitro* stability of the Cdc13/telomeric ssDNA complex. We found that the DNA exchange by Cdc13 occurs rapidly at physiological temperatures, requires telomeric repeat sequence DNA, and is affected by ssDNA length. Cdc13 truncations revealed that the low-affinity binding site (OB1), which is distal from the high-affinity binding site (OB3), is required for this intermolecular dynamic DNA exchange (DDE). We hypothesize that DDE by Cdc13 is the basis for how Cdc13 “moves” at telomeres to alternate between modes where it regulates telomerase activity and assists in telomere replication.

## INTRODUCTION

Telomeres are the nucleoprotein endcaps on linear eukaryotic chromosomes (1). Among their many functions, they address the end replication problem, end protection problem, and confer a chronological lifespan to cells. The structure of a telomere somewhat resembles a resected DNA double-strand break (DSB) because the repeat-sequence DNA comprising the bulk of the telomere ends in a single-stranded (ss)DNA tail on the G-rich strand, which is the substrate used by the enzyme telomerase to extend the telomere (2). However, there are many proteins competing for this G-overhang *in vivo*. In the budding yeast *Saccharomyces cerevisiae*, these factors include the general ssDNA binding protein (ssBP) RPA, the telomere-specific ssBP Cdc13 (3), and a number of DNA helicases (*e.g.*, Hrq1, Pif1, and Sgs1) (4–7). One outstanding topic in the field is how all of these factors interact and function on the same substrate to yield a homeostatic telomere length and support general telomere maintenance.

RPA functions in virtually all DNA transactions in a cell, where it stabilizes ssDNA generated during DNA replication, recombination, and repair by virtue of its high affinity for ssDNA (8). RPA is an evolutionarily conserved heterotrimer consisting of subunits named Rpa1-3 in *S. cerevisiae* (and commonly referred to by their approximate molecular weights: RPA70, RPA32, and RPA14, respectively). This ssBP contains six oligonucleotide binding (OB) domains that are vital for ssDNA binding and protein-protein interactions. RPA binds ssDNA with 5ʹ-3ʹ polarity (9,10) using different binding modes that are affected by the length of the substrate (11) and post-translational modifications of RPA subunits (11,12). These multiple ssDNA binding interfaces and binding modes make RPA-ssDNA complexes dynamic, with molecular modelling (13) and single-molecule biochemistry (14–17) demonstrating that, for instance, RPA can freely diffuse along ssDNA. At telomeres, RPA participates in DNA replication and plays roles in telomerase loading, DNA end protection, and DNA strand exchange (18–21). By analogy to its role at fission yeast telomeres and during G-rich mini-satellite maintenance in budding yeast (22,23), *S. cerevisiae* RPA may also serve to prevent the formation of DNA secondary structures on the G-strand to facilitate the binding of other telomeric factors and promote telomerase activity.

Cdc13 is another OB fold-containing ssBP in *S. cerevisiae* (Fig. 1). Extensive biochemical research confirms that Cdc13 is a central regulator of telomere length homeostasis and chromosome end protection as part of the Cdc13-Stn1-Ten1 (CST) complex (24–31). With Cdc13, Stn1, and Ten1 displaying similarities to Rpa1, Rpa2, and Rpa3, respectively (3), CST is often referred to as the telomere-specific RPA-like complex in *S. cerevisiae*. Regulation of CST complex formation is critical for the control of telomere extension, protection of telomeric ssDNA from nuclease degradation, and control of DNA polymerase alpha (Polα) activity (30,32–35). Regulation of Cdc13 is achieved in part by phosphorylation of Cdc13 and other components of the CST complex, affecting protein-protein interactions (32,36–38). During S-phase, Cdc13 and Stn1 phosphorylation causes Stn1 and Ten1 to dissociate from Cdc13 (39), allowing Cdc13 to directly interact with telomerase to begin telomere extension (31,40,41). As telomere extension proceeds, Cdc13 SQ/TQ sites are increasingly phosphorylated, and Cdc13 is also SUMOylated, while Stn1 is dephosphorylated, allowing the CST complex to reform (42). This enables Cdc13 and Stn1 to interact with Polα and the helicase Pif1 to fill in the telomere C-strand, completing telomere synthesis (25,43,44).

**Figure 1.**
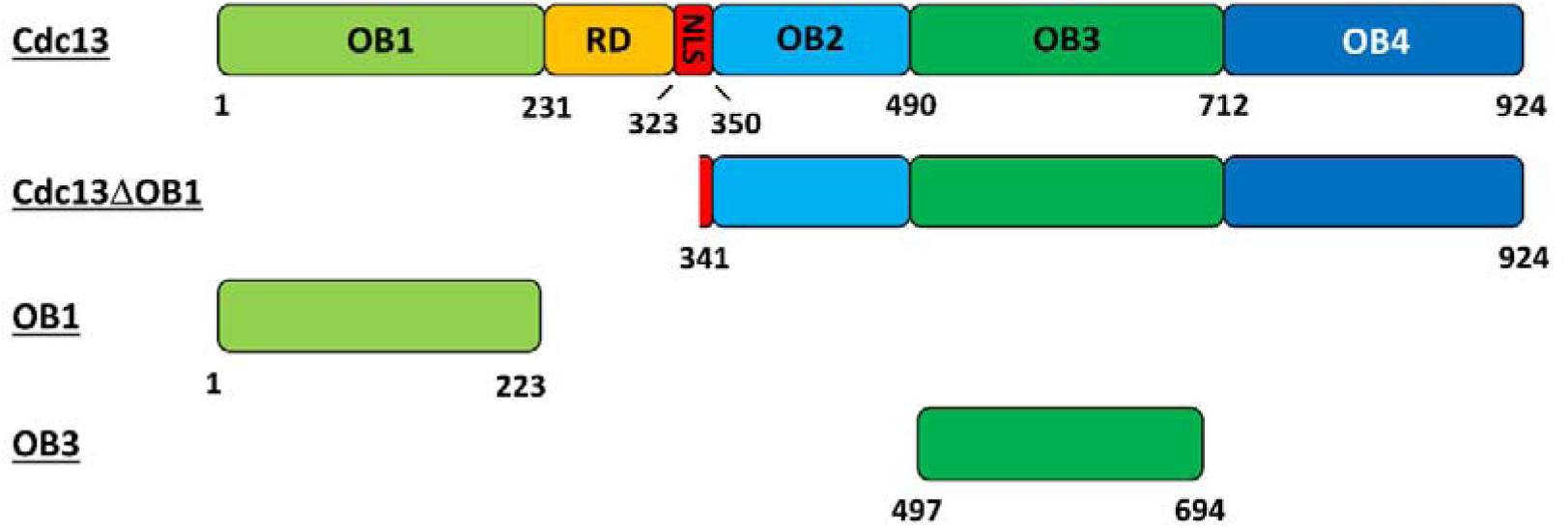
Domain schematics of recombinant proteins used in this work. The domains comprising full-length Cdc13 are shown to scale, with the numbers below the image indicating the amino acid number at the domain boundary. The domain schematics and sizes of the CDC13ΔOB1, OB1, and OB3 constructs are also shown. RD, telomerase recruitment domain; NLS, nuclear localization sequence.

Analyses of Cdc13 have identified four oligonucleotide/oligosaccharide binding (OB) folds (OB1-OB4), a telomerase recruitment domain (RD), and a nuclear localization signal (NLS) (Fig. 1) (30,45–48). ssDNA binding by Cdc13 has been rigorously investigated using wild-type protein, truncation mutants, and full-length mutants based on temperature-sensitive alleles, and the OB1 and OB3 domains have been shown to act as ssDNA binding domains (35,45,49–51). The full-length protein is reported to form an incredibly stable complex with telomeric ssDNA, displaying a 41-h half-life *in vitro* (52). As an isolated peptide, OB1 displays low-affinity for telomeric ssDNA (*K*_D_ ≈ 600 nM), and it requires long (≥37-43 nt) ssDNA substrates for binding (43,53). In contrast, OB3 in isolation has very high affinity (*K*_D_ ≈ 2 pM) for telomeric ssDNAs as small as 11 nt (47,52,54–57). OB1 and OB3 are connected by a flexible region (aa 231-350) that contains the Est1 binding sites (*i.e.*, the RD), the NLS, and the OB2 domain. A truncation mutant containing most of the predicted OB3 fold (aa 497-694) (Fig. 1) is sufficient for ssDNA binding *in vitro*, but *in vivo*, both OB2 and OB3 are required to bind to DNA in the nucleus because ablating OB2 disrupts the NLS (48).

Cdc13 appears to exist solely as a homodimer in both the cytoplasm and nucleus, and dimerization may play an important role in ssDNA binding (30,48,53). The L91R point mutation in the OB1 domain is known to abolish dimerization (43,53), but the OB2 and OB4 domains are also implicated in the formation of the Cdc13 homodimer (58) (Fig. S2). It has been demonstrated that mutations in OB2 lead to a loss of OB2-OB4 intramolecular interactions, but it is not entirely clear if they also disrupt intermolecular interactions (30,48,53). However, the *in vitro* DNA binding characteristics of the protein encoded by the *cdc13-1* allele (OB2 mutation) are essentially identical to those of wild-type Cdc13 (48).

One aspect of ssBP function at telomeres that has not been addressed is how proteins with incredibly tight binding affinities for telomeric ssDNA are removed after binding to complete replication and/or enable telomerase activity. It is possible that the DNA replication complex itself, with its associated Rrm3 helicase activity (59), could forcibly evict Cdc13 (60,61). However, other helicase-related mechanisms are also plausible. For instance, mutation of either the Pif1 or Hrq1 helicases increases the rates of gross chromosomal rearrangements (GCRs) that are preferentially healed by telomere addition (4), suggesting that these enzymes normally function as telomerase inhibitors. *In vitro*, Hrq1 and Pif1 directly regulate telomerase activity (6), and they may also be able to evict Cdc13 from telomeric ssDNA or the 3ʹ ssDNA caused by resection of DSBs to inhibit telomerase recruitment and thereby indirectly inhibit telomerase. Indeed, Pif1 can remove, destabilize, or block Cdc13 binding at short DSBs and offers a mechanism for protection of short DSBs from telomere addition (62–65).

Our research has focused on helicase-telomerase interactions and the roles they play in establishing telomere length homeostasis (6,66). The interaction of Pif1 with Est2 is known to inhibit the length of the 3ʹ overhang of telomeres *in vivo* (4,67). However, using *in vitro* telomerase assays, titration of Pif1 reveals a biphasic effect of Pif1 on telomerase activity (6). At low Pif1 concentrations, telomerase extension activity is increased, but an inhibition of telomerase activity occurs when Pif1 concentrations exceed 100 nM. Addition of Hrq1 along with Pif1 further decreases the level of telomerase activity, and this effect is independent of Hrq1 helicase activity (4,6). Hrq1 alone or in the presence of catalytically inactive Pif1 acts as an activator of telomerase activity (6). Similar experiments with the Sgs1 helicase, a RecQ family enzyme like Hrq1, show that it can also act as an activator of telomerase (5).

Cdc13 has also been reported to inhibit telomerase activity *in vitro* by limiting access of telomerase to 3ʹ telomeric ssDNA (41). Reports of Pif1’s ability to remove Cdc13 from telomeric DNA are mixed, with evidence presented that Pif1 both can and cannot remove Cdc13 from DNA dependent on the length or condensation state of the Cdc13-bound DNA (35,64,65). Furthermore, different groups have reported that Pif1 inhibits telomerase *in vivo* at all telomeres regardless of the length of the ssDNA overhang (68). Single-molecule analysis with immobilized telomere mimics shows that Pif1 preferentially removes telomerase from longer ssDNA. Several groups have reported that HO endonuclease-generated DSB overhangs with telomere dsDNA of ≥34 bp are insensitive to Pif1 activity, allowing increased telomerase extension (64). It is hypothesized that this represents a transition point between DSBs and telomeres, which is due to higher-affinity Cdc13 binding to these substrates. These results reinforce the view that there is a complex relationship between DNA helicases and other telomere binding proteins controlling telomere length and telomerase activity.

Here, we attempted to determine the effects of *S. cerevisiae* Cdc13 and RPA on Hrq1 and Pif1 helicase activity, with the goal of establishing *in vitro* telomerase activity assays that more closely mimic physiological conditions by including a more complete set of proteins associated with telomeres. Basic biochemical characterization of the ssBPs was performed, followed by experiments to determine if Hrq1 and/or Pif1 can remove Cdc13 or RPA from telomeric ssDNA. However, early experiments failed due to the collapse of the Cdc13-telomeric ssDNA complex, which should have been incredibly stable *in vitro* (47,52,55). We found that the unlabelled telomeric ssDNA used as a protein trap in the reactions induced a rapid exchange of Cdc13 from the labelled to unlabelled ssDNA. This dynamic DNA exchange (DDE) of Cdc13 between substrates was dependent on ssDNA length and sequence and only occurred at physiological temperatures. Truncation mutants of Cdc13 demonstrated that the two identified ssDNA binding domains, OB1 and OB3 (Fig. 1), are both required for DDE. We propose a model where DDE by Cdc13 allows for fast and efficient exchange between telomeres or along telomere ssDNA during extension by telomerase without full dissociation.

## MATERIAL AND METHODS

### Plasmids

The *Escherichia coli* vector for over-expression of the N-terminally His_6_-tagged Cdc13 OB3 domain (a.a. 497-694; Fig. 1A) (52) was provided by Deborah Wuttke. All other Cdc13 plasmids used for protein overexpression in this study were constructed in house using the pET28a vector and included an N-terminal His_6_-SUMO tag and a C-terminal 3xFLAG tag. Cloning details are available upon request. Untagged RPA was expressed from the pET11c-RPA, which encodes all three subunits of the heterotrimer (Rfa1-3), which was kindly provided by Dr. Ilya Finkelstein.

### Recombinant protein production

#### E. coli*-generated proteins*

*E. coli* strains Nico21 (DE3) pLysS or BL21-codon plus (New England BioLabs) were used for overexpression, and the purification procedure for His_6_-SUMO-tagged proteins has been previously reported (Nickens 2021). Briefly, cells were transformed with a pET28a-based protein expression plasmid and maintained on LB medium supplemented with 50 μg/mL kanamycin and 34 μg/mL chloramphenicol. Cell harbouring pCdc13-DBD (a.a. 497-694) were supplemented with 100 μg/mL ampicillin and 34 μg/mL chloramphenicol. Liquid cultures for protein overproduction were grown in LB or 2xYT medium with the same antibiotics listed above. Expression of the target protein was induced with 0.5 mM IPTG when the culture reached an absorbance of 0.8 at 600 nm, followed by overnight incubation at 16°C. Cells were collected via centrifugation, with average yields of 10 g of cell pellet per 4 L of culture. The cells were resuspended in 50 mL of K buffer (20 mM of KH_2_PO_4_ (pH 7.4), 500 mM KCl, 10% glycerol, 0.5 mM EDTA, 0.01% NP-40, and 0.2 mM β-ME) supplemented with a cocktail of protease inhibitors (aprotinin, chymostatin, leupeptin, and pepstatin A at 5 μg/mL and 1 mM phenyl-methyl-sulfonyl fluoride). Cells were disrupted by sonication for 9 min (15 s sonication, 30 s rest; 12 times), and lysates were clarified by ultracentrifugation (20,000 x g for 20 min) and incubated with 0.4 mL of His-Select nickel resin (Sigma) for 1 h at 4°C with gentle mixing. The resin was pelleted at 1000 g for 5 min and transferred to a gravity column. The matrix was washed twice with 10 mL K buffer containing 500 mM KCl, 0.1% NP-40, and 15 mM imidazole and then washed once with 10 mL of K buffer containing 500 mM KCl, 0.01% NP-40, and 15 mM imidazole. Protein was eluted with 2.5 mL of K buffer containing 200 mM imidazole. The eluate was then gently mixed with 300 μL of anti-FLAG-M2 resin (Sigma), incubated for 1.5 h or overnight at 4°C, and washed as above. The target protein was eluted twice by incubating the matrix with 0.5 mL of K buffer supplemented with 200 μg/mL FLAG peptide (Sigma) for 0.5 h, followed by a third elution step with no incubation. The eluate was filter-dialyzed into K buffer in an Ultracel-30K concentrator (Amicon) and further concentrated to ∼100 μL. Purified proteins were divided into 20-μL aliquots, frozen in a metal block submerged in dry ice, and stored at −80°C. Purity was assessed by SDS-PAGE and western blot analyses. This method was used to generate full-length Cdc13, the Cdc13ΔOB1, OB1 and OB3 truncations, and Pif1. Figure S1 shows images of all purified proteins separated by SDS-PAGE and visualized with Coomassie staining.

#### S. cerevisiae RPA purification

Untagged yeast RPA was purified as previously reported (69) with minor modifications. Cell pellets from 2 L of *E. coli* culture were thawed and resuspended in 50 mL resuspension buffer (20 mM Tris-HCl (pH 8.0), 500 mM NaCl, 5% glycerol, 0.01% NP-40, and 1 mM β-ME) supplemented with the protease inhibitor cocktail listed above, then lysed by sonication on ice for 10 min (10-s pulse on, 30 s off, 100% amplitude). The lysate was handled at 4°C for all subsequent steps. It was first clarified by centrifugation at 10,000 g for 30 min and loaded onto a 10-mL self-assembled Affi-Gel Blue gravity column (Bio-Rad). The column was sequentially washed with 50 mL resuspension buffer containing 0.3 M NaCl, 50 mL 0.5 M NaSCN, and 50 mL 1.5 M NaSCN. RPA eluted in the wash with 1.5 M NaSCN. The eluent was then applied to a 10-mL self-assembled HAP column (Bio-Rad), washed with 50 mL resuspension buffer, and eluted with 50 mL 100 mM potassium phosphate buffer (pH 8.0, with 0.01% NP-40 and 1 mM β-ME added). At this point, the RPA was 80% pure and concentrated to 1 mL using a Millipore Amicon UItra centricon (5 mL, 30 kDa cut-off) before application to a prepacked Superdex 200 10/300 GL gel filtration column (GE Healthcare) using resuspension buffer as the mobile phase. The fractions containing RPA were pooled, concentrated to 1 mL, and diluted to 100 mM NaCl by adding 4 mL resuspension buffer lacking NaCl. This material was then loaded onto a 5-mL prepacked MonoQ column (GE Healthcare) and eluted with a salt gradient from 100-500 mM NaCl. The purified RPA (Fig. S1) eluted between 200 and 230 mM NaCl, was slightly concentrated to 2-3 mg/mL, and stored in 10-μL aliquots at −80°C.

#### Insect cell-generated Cdc13

*Spodoptera frugiperda* Sf9 tissue culture was used to overexpress full-length, wild-type Cdc13 with a 5ʹ-SUMO-His_6_ tag and a 3ʹ-FLAG tag by subcloning this construct from the *E. coli* expression vector into pFastBac (Invitrogen). Bacmid DNA was generated by recombination in DH10bac cells. PCR screening was used to identify recombinants using primers MB1861 (CCCAGTCACGACGTTGTAAAACG) and MB1862 (AGCGGATAACAATTTCACACAGG). Cdc13-encoding baculovirus samples were generated by transfecting of 1 µg of bacmid DNA in 8 µL Cellfectin II reagent (Gibco) into 8×10 Sf9 cells in one well of a 6-well plate. Sf9 cultures were grown in SF900 media and infected with Cdc13-encoding baculovirus at a multiplicity of infection of 0.1. Cells were harvested by centrifugation after 3 days at 27°C with gentle aeration, and cell pellets were stored at −80°C. For protein purification, frozen cell pellets were thawed and resuspended in 50 mL of K buffer (20 mM of KH_2_PO_4_ (pH 7.4), 500 mM KCl, 10% glycerol, 0.5 mM EDTA, 0.01% NP-40, and 0.2 mM β-ME) supplemented with the protease inhibitor cocktail listed above and including DNase I at 20 µg/mL. Cells were lysed using a tissue homogenizer, and lysates were clarified by centrifugation (33,746 x g for 20 min) at 4°C. Clarified lysates were added to 50-mL conical tubes containing 2 mL of Ni resin (equilibrated in K buffer) per 25 mL of lysate and incubated for 1 h at 4°C with gentle rocking. The resin was gently pelleted, and the supernatant was removed before washing the resin three times with K buffer supplemented with 15 mM imidazole. Cdc13 was eluted from the resin by washing three times with 2 mL of K buffer supplemented with 200 mM imidazole. Eluates were pooled and mixed with 0.5 mL of anti-FLAG G1 affinity resin (GenScript) and incubated overnight in 15-mL conical tubes. The anti-FLAG resin was washed three times with 15 mL of K buffer, and Cdc13 was eluted by incubating the resin with 2 mL of K buffer supplemented with 200 µg/mL of 3x FLAG peptide (Sigma-Aldrich). Eluates were pooled and concentrated using Amicon Ultra 0.5 mL 100 k MW cutoff centrifuge filters (Millipore). Concentrated eluates were then applied to a 24-mL Superdex S200 increase 10/300 GL size exclusion column (GE Life Sciences). Fractions containing Cdc13 (Fig. S1) were pooled and concentrated to ∼100-200 µL, divided into 5-μL aliquots, frozen in liquid nitrogen, and stored at −80°C. Purity was assessed by SDS-PAGE and western blotting analyses. Hrq1, also generated in baculovirus-infected insect cells, was over-expressed and purified as described (70).

### Electrophoretic mobility shift assays (EMSAs)

Substrate oligonucleotides for all assays were purchased from Integrated DNA Technologies (IDT) and are listed in Supplementary Table S1. Binding reactions were performed in 1x binding buffer (25 mM HEPES (pH 8.0), 5% glycerol (w/v), 50 mM NaOAc, 150 mM NaCl, 7.5 mM MgCl_2_, and 0.01% Tween 20 (w/v)). Substrates were added to binding reactions to a final concentration of 2 or 2.5 nM as indicated. Binding reactions were incubated at 30°C for 30 min, then mixed with 6x non-fluorescent loading buffer (50 mM Tris (pH 8.0), 25% glycerol (w/v), and 0.025% Orange G) before separation on native 8%, 37.5:1 acrylamide:bisacrylamide gels in 1x Tris-glycine running buffer (25 mM Tris and 185 mM glycine (pH 8.8)). Gels were run at 100 V for 30-45 min and were imaged directly in their glass casting plates with a LICOR Odyssey DLx scanner and quantified using LICOR Image Studio software.

### Helicase assays

Fork substrates for helicase assays were constructed by incubating two partially complementary oligonucleotides (both at 1 μM) for annealing overnight at 37°C in a buffer consisting of 20 mM Tris-HCl (pH 8.0), 4% glycerol, 0.1 mM ethylene diaminetetraacetic acid, 40 μg/mL bovine serum albumin, 10 mM dithiothreitol, and 10 mM MgOAc (66). All DNA fork substrates used for helicase assays are listed in Table S1. We used either 5’ near infra-red (IR)-labelled probes (IDT) or 5’-^32^P-labelled probes to visualize the results. Radioactive oligonucleotides were ^32^P-labelled with T4 polynucleotide kinase and [γ P]-ATP under standard conditions. Radiolabelled [γ P]-ATP was purchased from PerkinElmer Life Sciences. Labelled oligonucleotides were separated from unincorporated label using G-50 microcolumns (GE Healthcare). One fork substrate was prepared with an IR 700 nm (IR700)-labelled oligonucleotide and an unlabelled oligonucleotide to create a fork with a poly(T)25 ssDNA and a Tel25G ssDNA and a 20 bp random dsDNA. A second fork substrate with dual Tel25G ssDNA and random dsDNA was prepared with a ^32^P-labeled oligonucleotide MB1170. Helicase reactions were performed at 30°C for 30 min in 1x EMSA binding buffer supplemented with ATP to a final concentration of 5 mM. Labelled fork substrates were added to a final concentration of 2 nM. Reactions were started by the addition of Pif1. Reactions were stopped by adding SDS and proteinase K to each reaction to a final concentration of 0.008% and 1.67 μg/mL, respectively, and incubated at 30°C for 30 min. Samples were then mixed with 6x non-fluorescent load buffer and placed on ice. Reaction products were separated on 10%, 19:1 acrylamide/bisacrylamide native gels using the same conditions as described above for EMSAs. Gels with radioactive label were dried, imaged, and quantified using a Typhoon 9500 scanner with ImageQuant software. Gels with IR labels were visualized using a LI-COR scanner, and images were quantified using Image Studio Lite version 5.2

### DDE assay

The DDE assay is essentially a modified EMSA. The labelled oligonucleotide substrate was mixed to a final concentration of 2 nM on ice with Cdc13 at 7.5 nM (3.75 nM dimer), unless otherwise noted, in a volume of 10 µL. These binding reactions were then incubated for 15 min at 30°C in 1x binding buffer. Competitor substrate, either unlabelled or conjugated to a different fluorescent label, was then titrated into the binding reactions and incubated for an additional 15 min at 30°C. Reaction products were separated on 7 or 8% native PAGE gels, imaged, and quantified as described above.

### Biolayer interferometry (BLI)

BLI was used to analyse protein association and dissociation kinetics from ssDNA substrates on a BLItz instrument (ForteBio). All BLI assays were performed in 1x binding buffer without glycerol at room temperature following the manufacturer’s protocols. Initial titration experiments were conducted to determine the concentrations of biotinylated Tel30G ssDNA probe and protein needed to saturate the streptavidin sensors. Based on titrations, biotinylated probe was bound to the sensor at 10 µM for 2 min. The association of 1 µM Cdc13 or 0.5 µM RPA to Tel30G was recorded for 2 min. To observe protein dissociation, the BLI sensor was moved to a large volume (1 mL) of EMSA binding buffer for 2 min (or longer as indicated), and DDE was observed by exposing the sensor to a 2- to 4-min incubation in the presence of 50 µM unlabelled competitor oligonucleotide. All BLI assays were performed at room temperature (∼23°C).

### Statistical analysis

Biological replicates (≥3) were performed for all reactions. The graphed data points are the means of these replicates, and the error bars are the standard deviation. All graphs were plotted using GraphPad Prism software, and where curves are fitted to the data, the details about curve fitting are provided in the figure legends.

## RESULTS

### *S. cerevisiae* ssBPs preferentially bind to telomeric ssDNA

In an effort to introduce additional telomere-binding proteins into *in vitro* telomerase assays (6), we first focused on *S. cerevisiae* ssBPs. To that end, we purified Cdc13 and RPA using standard methods and assessed their binding affinities to telomeric and poly(dT) ssDNAs (Fig. 2). As expected, Cdc13 bound more tightly to a 30-nt telomeric repeat sequence ssDNA substrate (Tel30G) ssDNA than to a poly(dT) 30mer (Poly(dT)30), with the concentration of protein required to bind 50% of the substrate (*k*_1/2_) equal to 3.7 ± 0.3 nM and 26 ± 3 nM, respectively (Fig. 2A). To our surprise, RPA also bound to Tel30G with higher affinity (2.4 ± 0.5 nM) than Poly(dT)30 (7.0 ± 0.5 nM) (Fig. 2B), though a similar phenomenon has also been reported for human RPA (71). Because we next wanted to determine if the ssBPs affect helicase unwinding of DNA forks, we also used EMSAs to measure the affinity of Cdc13 and RPA for a DNA fork substrate. These assays revealed that both Cdc13 (*k*_1/2_ = 3.3 ± 0.5 nM) and RPA (*k*_1/2_ = 2.5 ± 0.6 nM) bind forks with similar affinity to Tel30G under the *in vitro* conditions used here (Fig. 2A,B).

**Figure 2.**
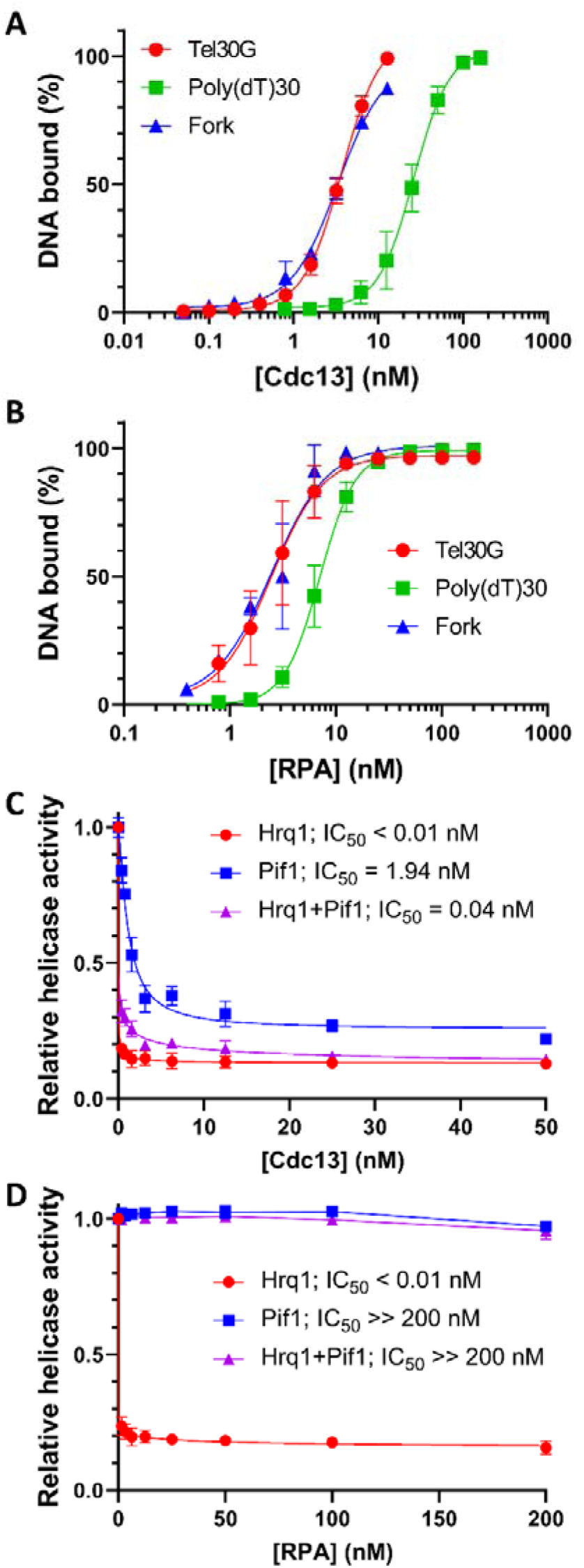
DNA binding by Cdc13 and RPA and its effect on Hrq1 and Pif1 helicase activity. Quantification of EMSA results for increasing concentrations of Cdc13 (A) or RPA (B) binding to 2 nM of the indicated substrates. The points are the average values from three independent experiments, and the error bars are the standard deviation (SD). C) Inhibition of DNA unwinding by Pif1 and/or Hrq1 helicase with Cdc13 prebound to a fork DNA substrate. Increasing concentrations of Cdc13 were added to helicase assays containing 2 nM telomeric arm fork substrate and 80 nM Hrq1, 30 nM Pif1 or both. In all three cases, Cdc13 was an effective helicase inhibitor. D) Increasing concentrations of RPA were added to helicase assays as described above. Here, RPA only inhibited Hrq1 alone. All points are the average values from ≥3 independent experiments, and the error bars represent the SD. The concentration of ssBP that inhibited helicase activity by 50% (IC_50_) is indicated on each graph. All reactions were incubated at 30°C, and reaction products were separated on gels at room temperature (∼23°C). In (A) and (B), the curves were fit with the Sigmoidal (4PL) equation, and in (C) and (D), the curves were fit with the Absolute IC50 equation (baseline constraint = 0) using GraphPad Prism.

### Inhibition of Hrq1 and Pif1 by ssBPs

We next determined the effect of the ssBPs on the ability of Hrq1 and Pif1 to unwind the fork substrate, which has a 20-nt mixed-sequence duplex region and 25-nt TG_1-3_ repeat sequence ssDNA arms. We found that Cdc13 inhibited the unwinding activity of both Hrq1 and Pif1, with Hrq1 being particularly susceptible (Fig. 2C). Similarly, the combination of both helicases was also inhibited by Cdc13. By holding the helicase concentration constant and titrating in Cdc13, we found that the concentration needed to inhibit unwinding by 50% (IC_50_) for Hrq1 was ∼0.01 nM, 1.94 nM for Pif1, and an intermediate value (0.04 nM) for the combination of Hrq1 and Pif1. These results indicate that Cdc13 is an effective inhibitor of Hrq1 and Pif1 *in vitro* helicase activity. Order-of-addition assays showed that adding the helicase first slightly reduced Cdc13 inhibition of unwinding activity, but the effect was small (data not shown).

Subsequently, we performed the same helicase inhibition assays with RPA. As with Cdc13, RPA was able to inhibit DNA unwinding by Hrq1 (IC_50_ < 0.01 nM; Fig. 2D). However, Pif1 helicase activity was largely unaffected, and the combination of Hrq1 and Pif1 displayed was likewise unaffected. These data correspond to recently published work demonstrating that human RPA does not act as a roadblock to *S. cerevisiae* Pif1 ssDNA translocation and is instead readily pushed along the nucleic acid by the helicase (6).

### Cdc13-Tel30G complexes are unstable *in vitro* in the presence of excess ssDNA

Although *S. cerevisiae* Pif1 is able to push human RPA into a DNA duplex to unwind 9 bp of dsDNA, it fails to unwind 18 bp by this method (16). In the assays above, our fork substrate contains a 20-bp duplex. Assuming that Hrq1 and Pif1 function similarly here, then if the ssBP is bound to the ssDNA arm of the fork along which the helicase translocates, the helicase must evict the ssBP to complete the unwinding reaction rather than pushing the ssBP along the ssDNA and through the dsDNA. Therefore, to determine if Hrq1 or Pif1 can remove ssBPs from DNA, we attempted to establish a gel-based assay to monitor the disruption of Cdc13-Tel30G complexes (Fig. S3). In these reactions, Cdc13 was prebound to the ssDNA, resulting in a gel shift relative to the unbound substrate, and then Hrq1 and/or Pif1 were added, along with excess unlabelled Tel30G to act as a protein trap. The Cdc13-telomeric ssDNA complex has a reported half-life of 41 h (52), so we expected to observe a loss of the Tel30G gel shift only if the helicases could evict Cdc13. However, we found that this happened equally well in control reactions lacking helicase but containing the unlabelled Tel30G (Fig. 3A). Indeed, time course experiments revealed that the addition of a 20-fold molar excess of unlabelled Tel30G primer caused ∼80% dissociation of preformed Cdc13-Tel30G complexes in 15 s (Fig. 3A). Therefore, despite using similar *in vitro* conditions as in the Figure 2A EMSAs and in published work in which the binding was stable (52), the Cdc13-Tel30G complex was unstable in the presence of unlabelled Tel30G trap in these experiments (Fig. 3A).

**Figure 3.**
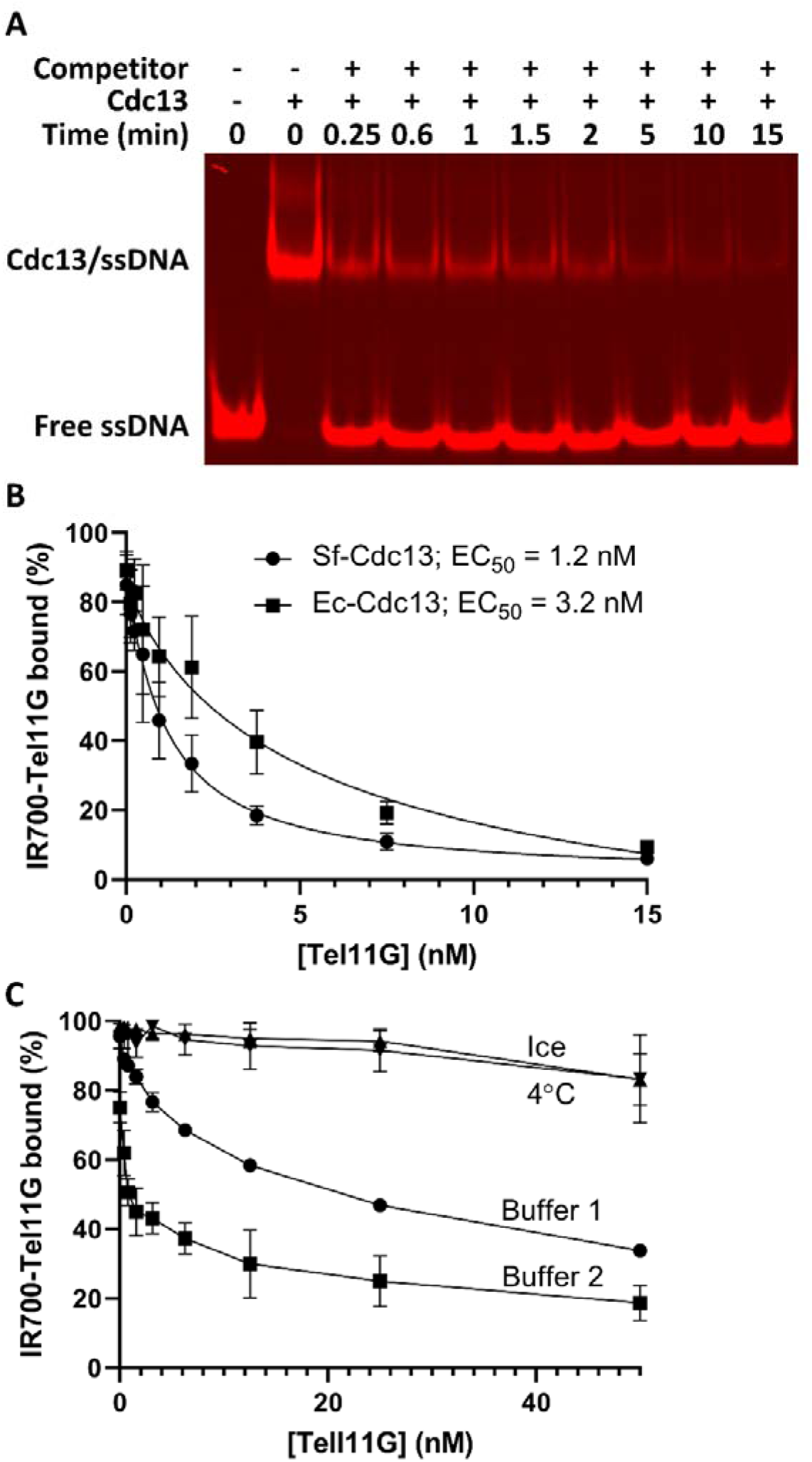
Cdc13-ssDNA complexes are unstable *in vitro* when competitor DNA is added at physiological temperatures. A) Representative gel image of the dissociation of Cdc13-Tel30G complexes over time when a molar excess of unlabelled Tel30G is added. The positions where free Tel30G and the protein-ssDNA complex migrate in the gel are noted. B) Protein-ssDNA complexes formed by 2 nM IR700-Tel11G and 3.25 nM recombinant Cdc13 dimer produced in either insect cells (Sf-Cdc13) or *E. coli* (Ec-Cdc13) both dissociate in the presence of unlabelled Tel11G competitor. The average values from three such experiments with SD are plotted, and the effective concentration of unlabelled Tel11G dissociating 50% of the protein-ssDNA complexes (EC_50_) is indicated. C) Dissociation experiments were performed as in (B), except for: 1) changing the incubation temperature from 30°C to either 4°C or incubation on ice, or 2) using the binding buffer from (52) (Buffer 2) rather than our typical binding buffer (Buffer 1). Only weak dissociation of Cdc13-Tel11G complexes was noted at high concentrations of unlabelled Tel11G competitor when the assays were performed at non-physiological temperatures, but effective dissociation was observed in both buffers. The average values from ≥3 independent experiments with their SD are plotted. All reactions were incubated at 30°C, unless otherwise noted, and reaction products were separated on gels at room temperature (∼23°C). In (B), the curves were fit with the Absolute IC50 equation (baseline constraint = 0) using GraphPad Prism.

Some previous work on Cdc13 used recombinant protein generated in *E. coli*, but here, we purified Cdc13 from baculovirus-infected insect cells. It is known that post-translational modifications (PTMs) affect Cdc13 activity *in vivo* (38), so our observation of unstable Cdc13-ssDNA binding could be due to PTMs that occur in insect cells but not *E. coli*. However, comparison of recombinant wild-type Cdc13 protein generated in both over-expression hosts revealed that they each readily dissociate from a labelled telomeric repeat sequence ssDNA 11mer (Tel11G) in the presence of even low concentrations of unlabelled competitor Tel11G (Fig. 3B), suggesting that PTMs do not affect this phenomenon.

Here, the shorter Tel11G substrate was used to more closely mimic the assays demonstrating the 41-h half-life for the Cdc13-Tel11G complex (52). However, we did use slightly different assay conditions (buffer and incubation temperature), and even seemingly simple buffer constituents can have large, unexpected effects on *in vitro* assays (72). Therefore, we repeated these assays using the published optimized DNA binding buffer at 30°C or at the previously used cooler incubation temperatures (52). As shown in Figure 3C, both buffer systems supported Cdc13-Tel11G dissociation, with the complex actually more readily dissociating in the optimized buffer (52). However, Cdc13 did not dissociate from the ssDNA when the reactions were incubated either at 4°C or on ice. Therefore, we were able to recapitulate the previously published stability of Cdc13-ssDNA complexes using psychrophilic conditions (≤4°C) but also uncovered a distinct Cdc13-ssDNA dissociation phenomenon that occurs at or near physiological temperatures (30°C).

### Dynamic DNA Exchange (DDE) by Cdc13

To investigate Cdc13-ssDNA dissociation further, we herein used a 30°C incubation temperature in all experiments (unless otherwise noted) and altered our assays to include Tel30G substrates conjugated to two different fluorophores: IR700 and IR800. By prebinding Cdc13 to IR800-Tel30G and then titrating in IR700-Tel30G, we could monitor the exchange of Cdc13 from one substrate to the other (Fig. 4A). These assays demonstrated that a 7.5-fold molar excess of IR700-Tel30G was sufficient to replace nearly all (94%) of the IR800-labeled substrate in the Cdc13-ssDNA complex (Fig. 4A,B). Control experiments switching the order of addition of the labelled substrates (*i.e.*, pre-binding to IR700-Tel30G) or using unlabelled competitor Tel30G yielded similar results (data not shown), demonstrating that the IR fluorophores did not affect the outcome of the assays. Similarly, to determine if this phenomenon was sequence dependent, IR700-labeled Poly(T)30 ssDNA was used as the competitor DNA. In this case, however, when Cdc13 was pre-bound to IR800-labeled Tel30G, the IR700-Poly(dT)30 completely failed to compete for binding, even at a 7.5-fold molar excess (Fig. 4C). Based on the observations above, we hypothesized that Cdc13 can dynamically exchange between ssDNA substrates, *i.e.,* it can ‘jump’ from one substrate to another in a process that is not simply driven by the off-rate of Cdc13 dissociating from its bound Tel30G ssDNA substrate. We termed this phenomenon DDE.

**Figure 4.**
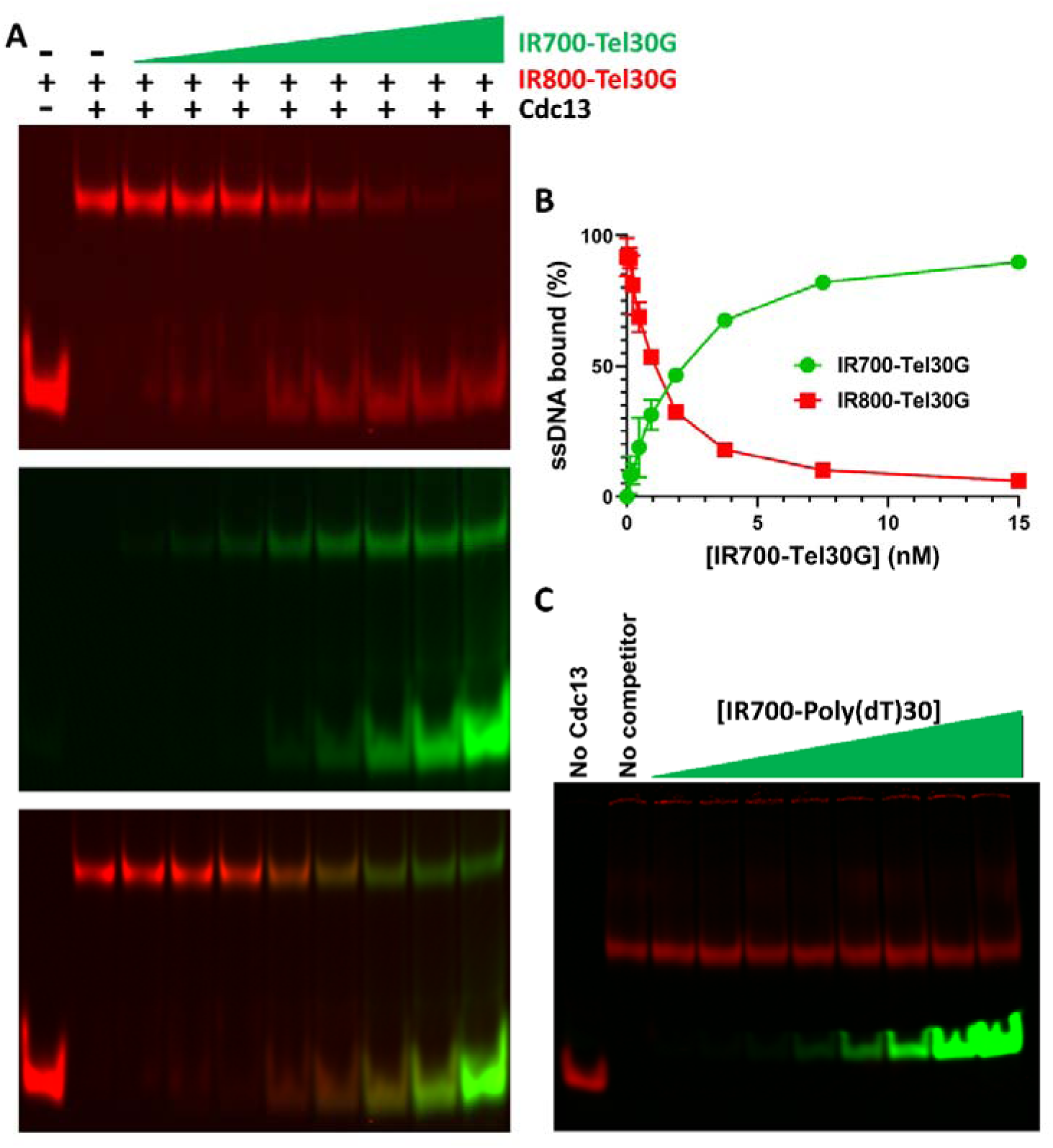
Dual-label dissociation assays demonstrate dynamic telomeric ssDNA strand exchange by Cdc13. A) Representative gel image of the dissociation of 3.75 nM Cdc13 dimer prebound to 2 nM IR800-Tel30G and then exposed to increasing concentrations of IR700-Tel30G as a secondary substrate. Top, IR800 channel; middle, IR700 channel; and bottom, merge. B) Quantification of triplicate assays performed as in (A), where the error bars are the SD. Even sub-stoichiometric concentrations of IR700-Tel30G effectively compete for Cdc13 binding. C) Representative gel image showing that increasing concentrations (0.12-15 nM) of IR800-Poly(dT)30 do not dissociate preformed complexes containing Cdc13 and IR700-Tel30G. All reactions were incubated at 30°C, and reaction products were separated on gels at room temperature (∼23°C).

Previous reports demonstrate that a telomeric 11mer ssDNA (Tel11G) is sufficient for high affinity binding by Cdc13 (47). To further investigate DDE by Cdc13, we next tested the effect of the length of the secondary ssDNA on the substrate exchange. We pre-bound Cdc13 to IR700-Tel30G and titrated in increasing concentrations of unlabelled Tel11G, 15G, 30G, and 50G ssDNAs (Table S1). In all cases, the secondary DNAs competed for Cdc13 binding, but the efficiency of DDE was inversely correlated to the length of the secondary substrate (Fig. 5A). The effective concentration of unlabelled ssDNA needed to reduce IR700-Tel30G binding by 50% (EC_50_) was equal to a 6.75-fold (13.5 nM) and 3.6-fold (7.2 nM) molar excess of Tel11G and Tel15G, respectively, relative to the pre-bound Tel30G (Fig. 5A). For Tel30G and Tel50G secondary substrates, the EC_50_ values were both sub-stoichiometric (0.89 and 0.96 nM, respectively; Fig. 5A) and similar, suggesting that the ssDNA length dependence of DDE efficiency hit a maximum at the same length as the primary IR700-labelled substrate.

**Figure 5.**
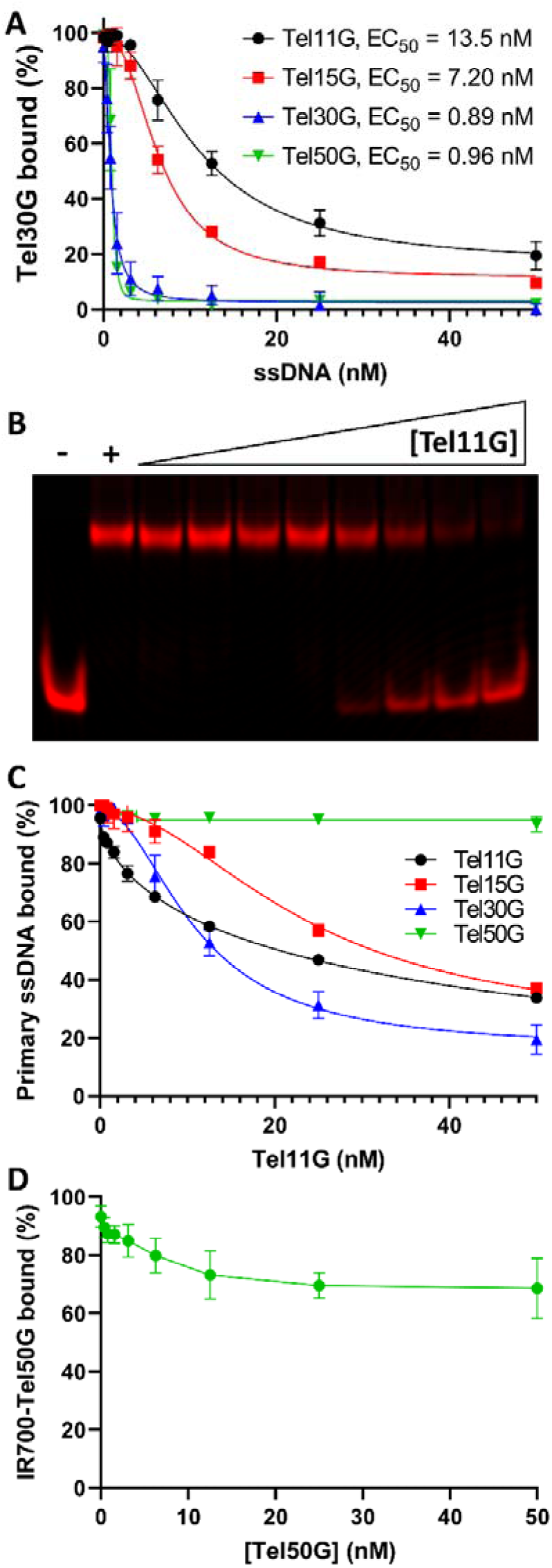
The effects of ssDNA length on DDE. A) Cdc13 (3.75 μM dimer) was prebound to 2 nM Tel30G, and then increasing concentrations of the indicated secondary ssDNA substrates were added. Sub-stoichiometric competition (indicative of DDE) was observed for the long substrates (Tel30G and Tel50G), but Tel15G and Tel11G demonstrated more modest abilities to compete for Cdc13 binding. The average values from three independent experiments are plotted, and the error bars are the SD. The EC_50_ of each secondary substrate is also noted. B) Representative DDE assay in which Cdc13 was prebound to IR700-Tel11G, and then unlabelled Tel11G (0.4-50 nM) was added as a secondary competitor substrate. The – and + symbols indicate reactions containing only IR700-Tel11G and IR700-Tel11G bound to Cdc13, respectively. C) Experiments were performed as in (A), but Cdc13 was prebound to each of the four substrates (Tel11G, 15G, 30G, and 50G) and then challenged with increasing concentrations of Tel11G as the secondary substrate. Overall, Tel11G was a poor competitor for Cdc13 binding (*i.e.*, no evidence of DDE), and completely failed to compete when Cdc13 was prebound to Tel50G. D) Cdc13 bound to Tel50G does not undergo DDE. Cdc13 was prebound to labelled Tel50G, and then increasing concentrations of unlabelled Tel50G were added as the secondary substrate. Even at 50 mM (25-fold molar excess), unlabelled Tel50G was a poor competitor. The average values from ≥3 independent experiments are plotted, and the error bars are the SD. All reactions were incubated at 30°C, and reaction products were separated on gels at room temperature (∼23°C). In (A) and (C), the curves were fit with the Absolute IC50 equation (baseline constraint = 0) using GraphPad Prism.

We next determined the effect of increasing the length of the prebound telomeric ssDNA on the efficiency of DDE using Tel11G as the secondary substrate. Tel11G was able to dissociate Cdc13 prebound to Tel11G, 15G, or 30G, but was unable to compete for binding when Cdc13 was prebound to IR700-Tel50G (Fig. 5B,C). In contrast, unlabelled Tel50G was able to disrupt the IR700-Tel50G:Cdc13 complex, albeit poorly, again demonstrating that longer ssDNAs are more efficient for Cdc13 DDE (Fig. 5D). Regardless, the relationship of the length of the prebound ssDNA to DDE efficiency was more complex when a short secondary substrate (Tel11G) was used, without a clear proportional relationship. Altogether, these assays demonstrate that Cdc13 is capable of inter-strand exchange between telomeric ssDNAs. Further, the sequence and length of both the bound and unbound ssDNAs impacts the efficiency of DDE.

### RPA also exhibits DDE

Having characterized the substrate requirements for DDE by Cdc13, we next sought to determine if RPA also exhibits this activity. Our assays demonstrated that DDE occurs when RPA is prebound to labelled Tel30G and exposed to unlabelled Tel30G as a secondary substrate, with EC_50_ = 5.44 nM (Fig. 6A). In contrast, Poly(dT)30 only weakly competed for RPA binding when RPA was prebound to Tel30G, even at a 50-fold molar excess (100 nM) of Poly(dT)30. However, when RPA was prebound to labelled Poly(dT)30 and exposed to unlabelled Poly(dT)30 as a competitor, then more effective competition occurred (EC_50_ = 15 nM; Fig. 6B). In contrast, though, Tel30G is a stronger competitor when RPA is prebound to Poly(dT)30 (EC_50_ = 5.1 nM). These data support the results from Figure 2B showing that telomeric ssDNA is a favoured substrate for RPA and that RPA can also undergo inter-strand exchange between telomeric ssDNA substrates.

**Figure 6.**
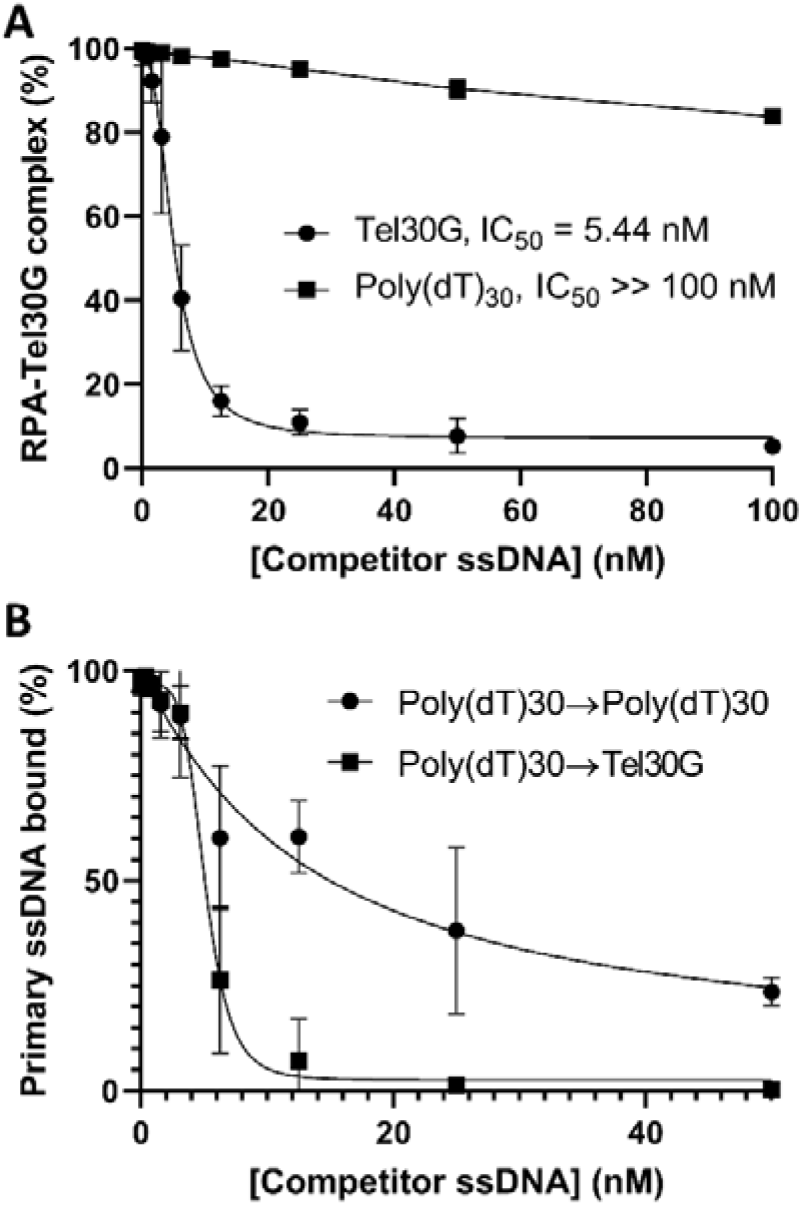
RPA also demonstrates DDE with telomeric ssDNA. A) RPA was prebound to Tel30G, and then increasing concentrations of unlabelled Tel30G or Poly(dT)30 were added to compete for binding. As with Cdc13, low concentrations of Tel30G efficiently dissociated RPA-Tel30G complexes, suggesting that DDE occurred, but Poly(dT)30 was a poor competitor. The average values from three independent experiments are plotted, and the error bars are the SD. The EC_50_ of each secondary substrate is also noted. (B) Experiments were performed as in (A), except RPA was prebound to Poly(dT)30 and then challenged with increasing concentrations of unlabelled Poly(dT)30 or Tel30G. Both secondary substrates could compete for RPA binding, but Tel30G was threefold more effective than Poly(dT)30 (EC_50_ = 5.1 nM *vs*. 15.1 nM). The average values from ≥3 independent experiments are plotted, and the error bars are the SD. All reactions were incubated at 30°C, and reaction products were separated on gels at room temperature (∼23°C). In (A) and (B), the curves were fit with the Absolute IC50 equation (baseline constraint = 0) using GraphPad Prism.

### DDE is dependent on intact Cdc13 protein

Moving forward, we decided to focus on the protein determinants of DDE by using a set of truncation mutants of Cdc13: the OB1 domain, the OB3 domain, and a construct lacking the OB1 domain (Cdc13ΔOB1) (Fig. 1). We started by characterizing OB3 because it is reported to be the primary DNA binding site of Cdc13 (52). Although it retained ssDNA binding activity, it failed to undergo DDE, with no binding competition observed at any concentration of secondary substrate added (Fig. 7A, S4). Therefore, the OB3 domain of Cdc13 is not sufficient for DDE alone, suggesting that another ssDNA binding site on Cdc13 is involved.

**Figure 7.**
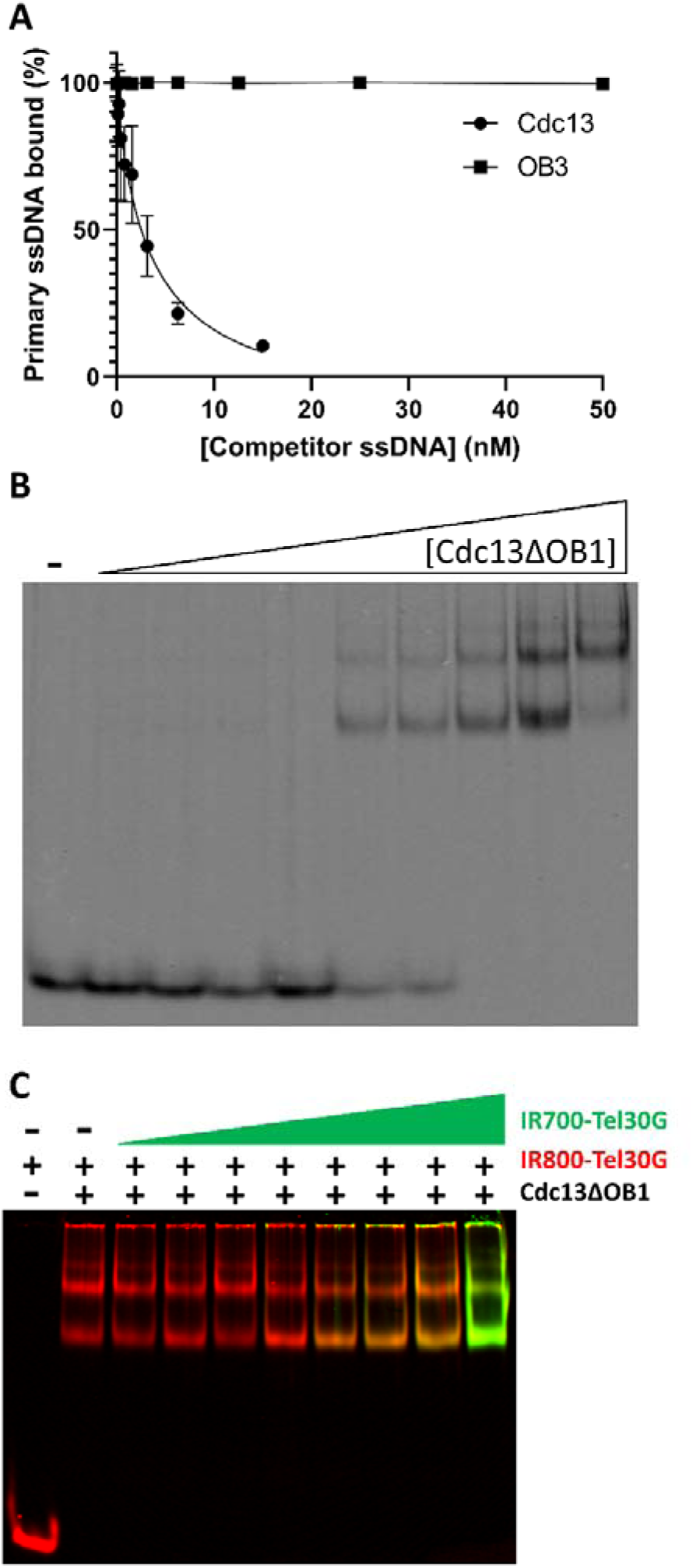
Only full-length Cdc13 undergoes DDE. A) OB3 was tested for DDE compared to full-length Cdc13, but preformed OB3-ssDNA complexes were stable in the presence of a molar excess of unlabelled competitor ssDNA. The average values from ≥3 independent experiments are plotted, and the error bars are the SD. The curves were fit with the Absolute IC50 equation (baseline constraint = 0) using GraphPad Prism. B) ssDNA binding by Cdc13ΔOB1. Representative EMSA gel of increasing concentrations of Cdc13ΔOB1 incubated with Tel30G. Multiple shifted species are evident, indicating more than one protein molecule bound to each ssDNA or *vice versa*. C) A representative DDE assay performed with Cdc13ΔOB1 prebound to IR800-Tel30G and then incubated with increasing concentrations of IR700-Tel30G. Instead of DDE or binding competition occurring, Cdc13ΔOB1 bound all of the additional substrate. All reactions were incubated at 30°C, and reaction products were separated on gels at room temperature (∼23°C).

It has been shown that the Cdc13 OB1 domain also has ssDNA binding activity (53), so we next investigated our OB1 domain construct. As reported, it did bind ssDNA, but unlike OB3, its binding affinity was poor (*k*_1/2_ = 830 nM), requiring micromolar amounts of protein to shift 100% of the Tel30G in an EMSA (Fig. S5A). Thus, to assess its ability to undergo DDE, we had to alter the standard assay by vastly increasing the concentration of the primary substrate to form the prebound OB1-Tel30G complex, as well as correspondingly increase the concentration of the competitor substrate. Under these conditions, we did observe competition for OB1 binding but only at a molar excess of competitor ssDNA (Fig. S5B).

To better compare activity to the full-length Cdc13 and OB3 assays but still focus on OB1, we instead used the Cdc13ΔOB1 truncation protein. In this case, Cdc13ΔOB1 bound to ssDNA with a comparable affinity to full-length Cdc13 and OB3, though the EMSAs displayed multiple shifted species, suggesting that Cdc13ΔOB1 may multimerize on ssDNA (Fig. 7B). The DDE assay with Cdc13ΔOB1 also revealed unexpected behaviour by the protein. Rather than exchange between substrates, Cdc13ΔOB1 bound all the additional ssDNA added to the reaction (Fig. 7C), with an average of 3.6 ssDNA strands bound by each Cdc13ΔOB1 dimer compared to an average of 1.2 ssDNAs per dimer of full-length Cdc13. These data suggest that in the context of the full-length Cdc13 protein, OB1 acts to restrict ssDNA binding by OB3.

### BLI analysis of DDE

As endpoint assays, EMSAs fail to provide high-resolution on- and off-rate ssDNA binding data, which we reasoned may be informative when studying a dynamic process like the DDE observed here. Therefore, we turned to BLI technology to monitor Cdc13-Tel30G binding and DDE in real time (73). For these assays, biotinylated Tel30G ssDNA was immobilized on streptavidin sensors, which were then dipped into protein solutions to initiate ssDNA binding. The sensors were then shifted to a large volume of binding buffer lacking protein to allow dissociation of the protein-ssDNA complex to occur, and finally, the probes were moved into a solution containing free Tel30G to allow DDE to occur. Representative examples of this workflow are shown in Figure S6, and one caveat of these experiments is that they were performed at room temperature (∼23°C).

Using this technique, we found that Cdc13 rapidly associated with the immobilized Tel30G substrate (Fig. 8A) but displayed no dissociation from the ssDNA (neglecting the first 30 s of dissociation data, described below in the Discussion) (Fig. 8B). We also longitudinally followed this dissociation step for up to 24 h and likewise observed no appreciable dissociation (data not shown). Therefore, as previously reported (52), our Cdc13-telomeric ssDNA complexes were incredibly stable *in vitro* in the absence of a secondary ssDNA substrate to bind to. However, movement of the Cdc13-Tel30G complexes to buffer containing free Tel30G resulted in loss of Cdc13 binding to the immobilized Tel30G (Fig. 8C) as observed in gel-based DDE assays (Fig. 3-5).

**Figure 8.**
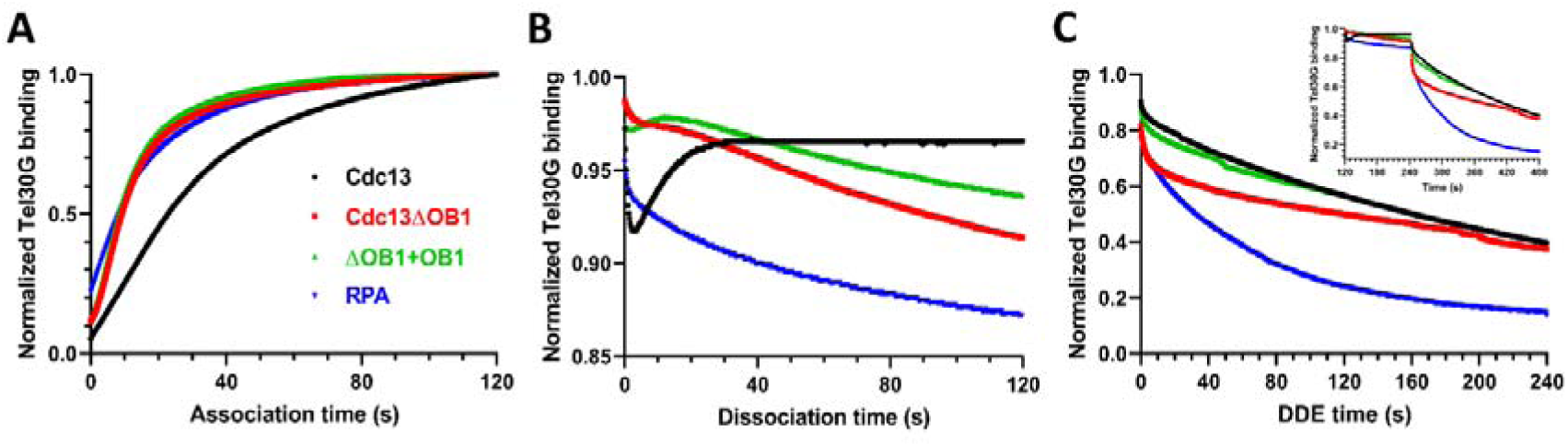
BLI analysis of Cdc13 and RPA binding to telomeric ssDNA. A) The association of Cdc13, Cdc13ΔOB1, an equimolar mix of Cdc13ΔOB1 and OB1 (Cdc13ΔOB1+OB1), and RPA with Tel30G was monitored for 2 min. B) Then, the protein-ssDNA complexes were diluted into a large volume of buffer, and dissociation of the proteins from Tel30G was monitored for 2 min. C) Finally, the remaining protein-ssDNA complexes were exposed to free Tel30G in solution, driving DDE. The inset shows the contiguous data from the dissociation and DDE steps to better visualize the changes in the rates of protein dissociation from the BLI sensor. The curves shown in all plots are comprised of data points collected every 0.2 s and are representative of ≥ 3 independent experiments. All BLI experiments were performed at room temperature (∼23°C).

BLI experiments were next performed with Cdc13ΔOB1. This truncation mutant bound to immobilized Tel30G with a faster association rate than Cdc13 (Fig. 8A), but unlike the full-length protein, Cdc13ΔOB1 dissociated from Tel30G when the protein-ssDNA complexes were diluted into buffer in the absence of free ssDNA (Fig. 8B). In the presence of free Tel30G, however, the off-rate increased (Fig. 8C), which differs from the results in our gel-based DDE assay showing super-stoichiometric binding of telomeric ssDNA by Cdc13ΔOB1 (Fig. 7C). This discrepancy could be due to the locally high concentration of ssDNA on the BLI sensor tip saturating the Cdc13ΔOB1 binding sites and/or the overall higher concentration (50 µM) of free Tel30G used in this assay.

Regardless, we sought to further investigate this by characterizing the OB1 construct. Here, it failed to robustly bind to Tel30G (data not shown), which is consistent with our EMSAs and reports that OB1 requires 43 nt of ssDNA for binding (53). Subsequently, we instead added it in *trans* to Cdc13ΔOB1 to determine if it could recapitulate the BLI profile of full-length Cdc13. Although this strategy had little effect on association with Tel30G compared to Cdc13ΔOB1 alone (Fig. 8A), it partially reproduced full-length Cdc13’s initial dip and rebound in binding during the dissociation phase of the reaction (Fig. 8B) and yielded a DDE curve more closely resembling that of full-length Cdc13 than Cdc13ΔOB1 (Fig. 8C).

Finally, we also characterized RPA-Tel30G binding using BLI. It readily bound Tel30G, with an association rate like that of Cdc13ΔOB1 (Fig. 8A), and RPA likewise exhibited significant dissociation in the absence of free ssDNA (Fig. 8B). In the presence of excess Tel30G, the off-rate increased (Fig. 8C), suggesting that DDE was occurring and corresponding to our gel-based DDE assays with RPA (Fig. 6).

## DISCUSSION

In this study, we used various *in vitro* approaches to characterize a novel phenomenon displayed by Cdc13 when interacting with telomeric repeat sequence ssDNA, which we term DDE. In contrast to the stable, high-affinity Cdc13-ssDNA binding that has been reported, we found that this binding activity is dynamic, with Cdc13 rapidly exchanging between substrates (Fig. 3A). However, DDE only occurred at physiological temperatures (Fig. 3C), which is a difference between our *in vitro* assays and much of the Cdc13 biochemistry that has been performed at 4°C or on ice in the past (47,52,74). It should be noted that Chib *et al.* (2023) recently reported a 15 nM *K*_D_ for Cdc13-Tel11G binding at 25°C (65), which is comparable to our result of *k*_1/2_ = 3.7 nM for Tel30G-Cdc13 binding at 30°C (Fig. 2A). DDE is both ssDNA sequence and length dependent, with Poly(dT) ssDNA not acting as a DDE substrate (Fig. 4C) and only telomeric substrates between 15 and 50 nt supporting the exchange (Fig. 5). From the perspective of the protein, both known ssDNA binding sites in Cdc13 (OB1 and OB3 (Fig. 1)) are necessary for DDE (Fig. 7), and recombinant Cdc13 produced in two different expression hosts is competent for DDE (Fig. 3B), arguing against an essential role for PTMs in the phenomenon. The general ssBP in *S. cerevisiae*, RPA, also displays DDE (Fig. 6, 8), though with the same telomeric repeat sequence dependency. Other ssBPs also display dynamic ssDNA binding (*e.g.*, *E. coli* SSB (75)), suggesting that this could be a general property of proteins that bind to ssDNA. The sequence specificity noted here indicates that DDE may also rely on a secondary structure that is formed by the telomeric ssDNA, though there is some discrepancy regarding the ability of *S. cerevisiae* (TG_1-3_)_n_ sequences to form stable secondary structures, especially in the presence of ssBPs (76,77). All of this is discussed in further detail below, especially regarding potential interactions with telomere-resident helicases and in telomere biology.

### DNA binding by Cdc13

Structural and biochemical analyses have identified the OB1 and OB3 domains of Cdc13 as the ssDNA binding domains in *S. cerevisiae* (30,58,74,78,79). These sites are distinguished by a wide range of affinities for ssDNA, with OB1 as a low affinity binding domain for DNA ≥ 43 nt, and OB3 as the high affinity binding domain that can bind very short (≥ 8 nt) ssDNAs (47,80). As an isolated domain, OB3 has a 4.5-fold higher ssDNA binding affinity than full-length Cdc13, leading to the conclusion that OB3 is <u>the</u> DNA binding domain (47,52). Published *K*_D_ values for full-length Cdc13 binding to Tel11G ssDNA are in the low picomolar range under psychrophilic *in vitro* conditions (47,52,74). Our binding assays were performed under different conditions, including temperature (30°C), and yielded an average *k*_1/2_ = 3.70 nM using Tel30G (Fig. 2A). This is comparable to the *K*_D_ of 5.2 nM reported for *Kluyveromyces lactis* Cdc13 from binding assays performed at room temperature (30). Regardless of ssDNA binding conditions, however, the Cdc13-ssDNA interaction is still high affinity.

As separate recombinant proteins, the OB1 domain of Cdc13 and Cdc13ΔOB1 displayed altered binding to telomeric ssDNA compared to full-length Cdc13 (Fig. 7,8). As previously reported, OB1 binds with low affinity to Tel50G DNA (*K*_D_ = 830 nM) and failed to bind to Tel30G (53). However, the Cdc13ΔOB1 truncation efficiently bound Tel30G (*k*_1/2_ = 0.724 ± 0.075 nM) (Fig. 7B), with a faster association rate than Cdc13, appreciable dissociation in the absence of a secondary substrate, and faster initial rate of DDE (Fig. 8). Despite the higher dissociation rate observed by BLI, Cdc13ΔOB1 also bound threefold more ssDNA than full-length Cdc13 – three-to-four Tel30Gs per Cdc13ΔOB1 dimer *vs.* one Tel30G per Cdc13 dimer – suggesting that the OB1 domain restricts the number of ssDNA molecules that can be bound by the OB3 domains in Cdc13 dimers (Fig. 7C). This was an unexpected result because Cdc13 dimers have four potential DNA binding sites, while Cdc13ΔOB1 has only two identified ssDNA binding domains (Fig. 1). These results have implications for understanding the functions of Cdc13 at telomeres and DSBs in TG-rich chromosomal regions and the interaction of the Hrq1 and Pif1 helicases with Cdc13 at these loci (4,81).

### DNA helicases and Cdc13

Despite the weaker ssDNA binding affinity that we report at physiological temperatures, Cdc13 is still a tenacious telomeric ssBP, which raises the question of how Cdc13 is removed from ssDNA to complete replication at chromosome ends. Possibilities include the replisome being able to remove a Cdc13 roadblock, perhaps aided by the PIF1 family helicase Rrm3 (7), or the Hrq1 and Pif1 DNA helicases (alone or in combination) displacing Cdc13 (35,82,83). Results from our helicase assays using fork substrates with telomeric G-strand arms demonstrate that Cdc13 inhibits DNA unwinding by both Hrq1 and Pif1 *in vitro*, with IC_50_s < 2 nM (Fig. 2C). In contrast, RPA failed to inhibit Pif1 and Hrq1+Pif1 helicase activity on the same fork substrate (Fig. 2D). This is unsurprising given recent single-molecule results showing that *S. cerevisiae* Pif1 can freely push human RPA on ssDNA (16). However, like Cdc13, we found that RPA potently inhibited Hrq1 helicase activity on the telomeric fork (Fig. 2D), despite telomeric ssDNA being a preferred substrate of Hrq1 (84,85). These data indicate that RPA can be moved or removed from telomeric ssDNA when Pif1 is present, but Cdc13 is more problematic, even with the synergistic activity of Hrq1 and Pif1 (6). This helicase inhibition difference between the ssBPs, which otherwise have similar binding affinities for telomeric ssDNA and the telomeric fork substrate, may simply come down to the low level of Cdc13 dissociation from telomeric ssDNA in the absence of a secondary telomeric substrate to bind to, while RPA readily dissociates under the same conditions (Fig. 8B).

### Cdc13 displacement from telomeric DNA

Removing Cdc13 from DNA ends by helicases could be important for many of the steps of telomere biology such as Okazaki fragment removal, C-strand fill in, and DNA end protection (2). One goal of our research is to understand the mechanism of how the CST complex dissociates and reforms on telomere DNA and the roles that Hrq1 and/or Pif1 might play in these processes. To approach this question, we used a modified EMSA that was designed to test whether Hrq1, Pif1, or the combination thereof could displace Cdc13 bound to telomeric ssDNA (Fig. S3). Early experiments failed due to a collapse of the Cdc13:ssDNA complex, despite the tight *k*_1/2_ that we found (Fig. 2A) and the reported stability of this nucleoprotein complex (52). Subsequent troubleshooting demonstrated that the presence of unlabelled or differentially labelled secondary telomeric ssDNA caused a fast dissociation or exchange event at physiological temperatures (Fig. 3), which we subsequently characterized as DDE.

A stated above, DDE is not simple dissociation of Cdc13 from ssDNA and is not driven by its off rate, which is miniscule in the absence of competitor ssDNA at both 4°C and room temperature (Fig. 3C, 8B). Indeed, Cdc13 undergoing DDE does not behave like a typical DNA binding protein in a binding competition assay where a large molar excess of competitor substrate is required. Here, exchange of Cdc13 from one Tel30G molecule to another was evident at sub-stoichiometric amounts of secondary substrate, with 50% DDE occurring at a 1:1 ratio of unlabelled-to-labelled (or IR700- to IR800-labelled) Tel30G (Fig. 4).

While telomeric sequence supports DDE and poly(dT) does not (Fig. 4), length is also critical (Fig. 5). DDE assays showed that when Cdc13 was prebound to Tel30G, Tel15G and especially Tel11G behaved more like competitors than DDE substrates because a molar excess was required to disrupt Cdc13-Tel30G binding (Fig. 5A). Similarly, Tel11G was a poor secondary substrate when Cdc13 was prebound to Tel11G, 15G, or 30G, only displaying competition at high concentrations and no DDE (Fig. 5B,C). With Cdc13 prebound to Tel50G, Tel11G failed to compete at all, even at a 25-fold molar excess, and unlabelled Tel50G was likewise a poor competitor when Cdc13 was prebound to labelled Tel50G (Fig. 5D), suggesting that binding to long stretches of telomeric ssDNA inhibits DDE by Cdc13. This feature offers a mechanism for Cdc13 to unilaterally exchange from short to long telomeric substrates, perhaps allowing Cdc13 to move toward the 3ʹ ssDNA end of a newly extended telomere.

Regarding the protein determinants of DDE, we found that the high-affinity ssDNA binding domain of Cdc13 (OB3) lacks DDE (Fig. 7A), and removal of the low-affinity site (OB1) in the context of the Cdc13ΔOB1 truncation yielded super-stoichiometric ssDNA binding (Fig. 7C). This precluded the observation of DDE at low ssDNA concentrations, but it appears to occur at higher concentrations in BLI assays (Fig. 8C). Taken together, these data indicate that both OB1 and OB3 domains control ssDNA binding and DDE by Cdc13. The region between these domains in the protein is predicted to be largely unstructured (with the exception of OB2) (Fig. S2), which is supported by partial crystal structures of Cdc13 (30,43,58). Therefore, we hypothesize that some portion of the disordered region between OB1 and OB3 is required for DDE, perhaps to communicate the ssDNA binding state of one domain to the other or to allow sufficient conformational dynamics to enable exchange between ssDNAs. This region also contains known sites of protein-protein interactions (*i.e.*, the RD and NLS (Fig. 1)), so it is tempting to speculate that other proteins that interact with Cdc13 in the nucleus may affect its DNA binding and DDE *in vivo*. However, it cannot be ruled out at this time that other regions of Cdc13 are involved in DDE, and future experiments will further investigate this phenomenon (86).

### Cdc13 and BLI

It should be noted that BLI observation of full-length Cdc13 dissociation from immobilized Tel30G in the absence of added free secondary substrate ssDNA was complicated by a sudden drop in the apparent size of the Cdc13-Tel30G complex after placing the sensor into a large volume of binding buffer to allow dissociation to occur (Fig. 8B). This decrease in the size of the complex rebounds in approximately 30 s to the original size observed during association. We have no explanation for this observation but hypothesize that it could be due to local dissociation of Cdc13 from one Tel30G molecule and reassociation with the same or a different Tel30G substrate on the molecularly crowded sensor tip. Full dissociation, rather than local dissociation, would result in a decreasing signal that does not rebound. Alternatively, Cdc13 may somehow condense the ssDNA to a shorter length upon the shift into buffer, with this condensation relaxing in ∼30 s. It has been reported that telomeric ssDNA can undergo a condensation or size reduction upon Cdc13 binding (35). This report is based on experiments using duplex telomeric DNA with relatively short 3ʹ ssDNA overhangs and there is no mention of the size recovering, but Tel30G condensation and subsequent decondensation could account for the dip and recovery in our BLI signal (Fig. 8B).

Regardless, we do not believe that this is simply a mechanical effect of moving the sensors between buffers because it does not occur between other steps in the assay that are performed likewise. Similarly, it is Cdc13 dependent, because we do not observe this dip with the other proteins tested, *e.g.*, RPA (Fig. 8B). Indeed, we can only come close to reproducing it by mimicking full-length Cdc13 with Cdc13ΔOB1+OB1.

### Model for Cdc13 DDE *in vivo*

Cdc13 is clearly a central regulator of DNA end protection and telomerase activity in budding yeast (29–31,87,88). Coordination of the binding to and dissociation of Ten1/Stn1 from Cdc13 is critical to control telomerase initiation, regulate subsequent telomerase activity, and interact with Polα to complete C-strand DNA synthesis (43,89). At this time, regulation of Cdc13 activities appears to be controlled by a complex set of PTMs, primarily phosphorylation events, occurring on Cdc13, Stn1, and Pif1 (32,38,90,91). Other modifications such as SUMOylation of Cdc13 and acetylation of Pif1 offer additional regulatory mechanisms (42,92,93). Cdc13 also plays a role in telomere addition at DSBs where the interaction of Pif1 and/or Hrq1 may be crucial for removing Cdc13 to avoid telomere addition to DSBs (4). It is unclear how PTMs may affect DDE by Cdc13 at this time.

One important question raised by the literature is how a protein with tight ssDNA binding affinity and a dissociation half-life of 41 h (52) removed from DNA in an organism with a 90-min doubling time. Some mechanism must exist to increase the rate of dissociation as telomere length increases *in vivo*, or Cdc13 would be trapped at its original DNA binding site. To date, the primary mechanism proposed for removal of Cdc13 by helicase activity is somewhat contradictory. One group reports that telomeres and DSBs become insensitive to Pif1 inhibition of telomerase when telomeric DNA is ≥ 34 bp, ostensibly due to Cdc13 binding (64). In contrast, another group reports that Pif1 fails to remove Cdc13 when bound to 15 nt of telomeric DNA in a low Brownian motion state but could remove Cdc13 in higher Brownian motion states on the same substrate (35). An additional complication is that Cdc13 retention in the nucleus is dependent upon binding to DNA. It has been proposed that Cd13 dissociation from DNA and subsequent transport out of the nucleus could be a mechanism for controlling telomere addition at DSBs (48). DDE provides an explanation for how Cdc13 can move on and between telomeres without direct dissociation from the DNA.

Our results show that Cdc13 is a potent inhibitor of Pif1 helicase activity (Fig. 2C), but our attempts to measure dissociation of Cdc13 by Pif1 in bulk gel-based assays failed due to the instability of the Cdc13-ssDNA complexes in the presence of unlabelled ssDNA as a protein trap (Fig. 3A). Our subsequent investigation of this DDE demonstrated that Cdc13 dynamically undergoes inter-strand transfer from one telomeric ssDNA to another. Due to the conformational flexibility of ssDNA, we hypothesize that Cdc13 may also be able to undergo intra-strand transfer from its original position to a distal open binding site of telomeric DNA at very low ratios of bound to unbound telomere DNA. We propose a model where the combination of Pif1 (±Hrq1) helicase activity and DDE allows Cdc13 to dissociate from telomeric ssDNA, though our results also support the conclusion that DDE alone may be able to account for Cdc13 removal from a telomere without invoking a helicase (Fig. 9).

**Figure 9.**
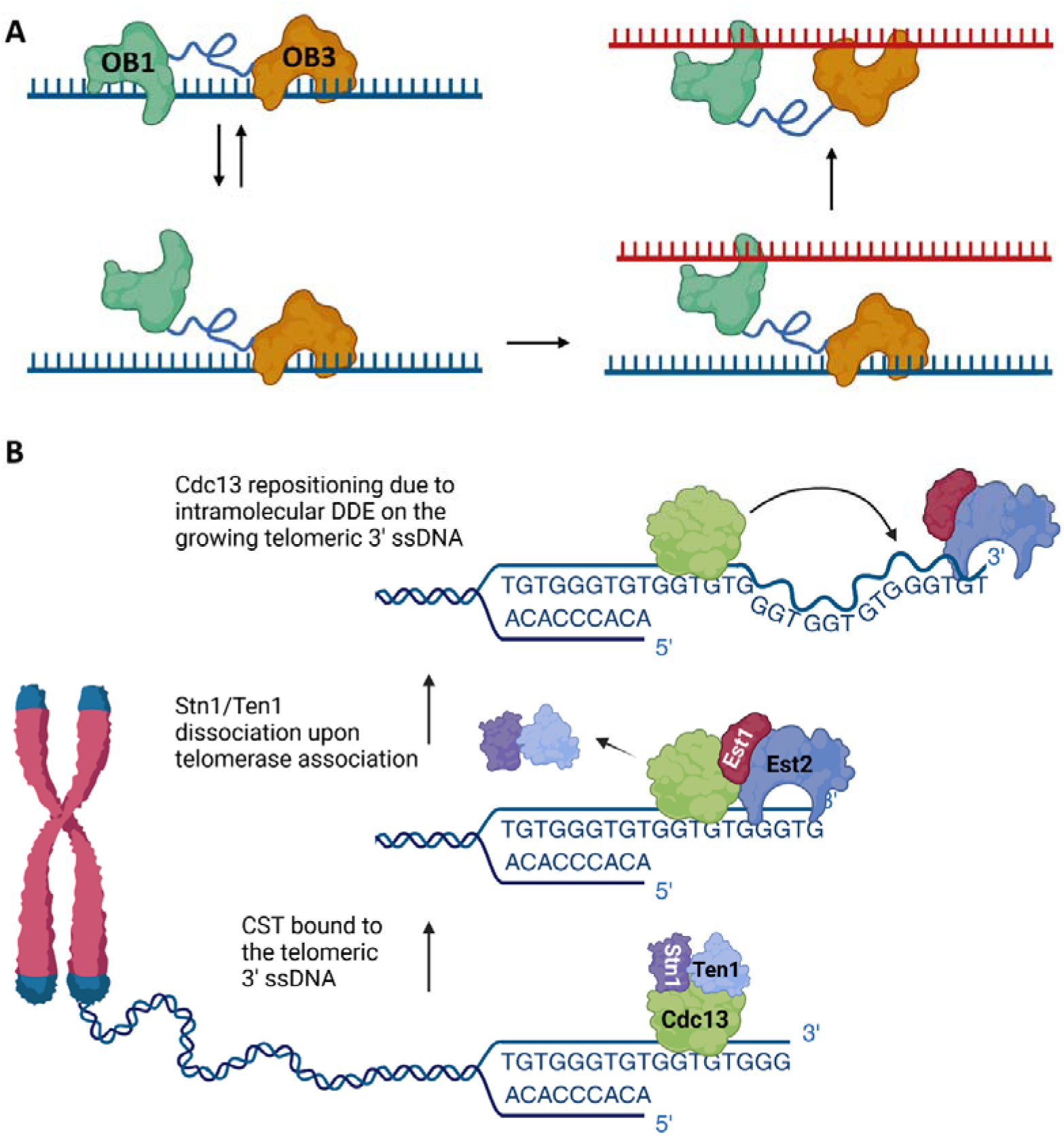
Model of DDE by Cdc13. A) When bound to ssDNA, Cdc13 may exist in an equilibrium between both binding sites (OB1 and OB3) being engaged with ssDNA and the low-affinity site (OB1) dissociating from ssDNA while the high-affinity site (OB3) remains bound. In the presence of a second ssDNA, OB1 can bind this substrate and communicate its binding state to OB3, which then allows OB3 to dissociate from the original substrate and bind to the second ssDNA. Here, Cdc13 is shown as a monomer for simplicity, but this general model holds true for two OB1 domains in a dimer binding a secondary substrate and communicating their ssDNA binding state to their cognate OB3s. B) A hypothetical model for intramolecular DDE by Cdc13 at an elongating telomere. As part of the CST complex, Cdc13 binds the 3ʹ telomeric ssDNA overhang. Upon recruitment of telomerase via Cdc13-Est1 interaction, Stn1 and Ten1 dissociate, and telomere lengthening begins. The telomeric ssDNA is conformationally flexible, which brings a distal binding site into proximity of Cdc13, enabling movement toward the 3ʹ end by DDE. In both (A) and (B), the activity of the Hrq1 and/or Pif1 helicases (not shown) may increase the off-rate of the OB1 domain or Cdc13 as a whole to facilitate DDE.

The mechanism for DDE is currently unknown, but we propose that when Cdc13 is bound to telomeric ssDNA, the high-affinity binding domain (OB3) remains associated, while the low-affinity domain (OB1) is either unbound or alternates between a DNA-bound and -unbound state (Fig. 9A). In the presence another unbound ssDNA binding site of sufficient length (either on another telomere or intramolecularly at a distal site), OB1 can bind to this secondary site. This association is transmitted through the linker between OB1 and OB3, or in *trans* across the Cdc13 dimer interface, altering the affinity of Cdc13 for the primary substrate and allowing high-affinity binding to the secondary substrate. This would lead to a ratchet-type movement, so Cdc13 remains bound to ssDNA even as it is exchanging strands.

It has been widely demonstrated in the literature that excess nucleic acids can facilitate the exchange of binding proteins by so-called direct transfer (see (94) and references therein), though a demonstration of biological relevance is still lacking. While we have only demonstrated inter-strand transfer by Cdc13, we hypothesize that intra-strand DDE on newly synthesized telomere DNA would increase the local concentration of free telomeric ssDNA, allowing Cdc13 to move toward the new 3ʹ ends (Fig. 9B). Other possible scenarios where DDE could come into play are Cdc13 movement to DSBs with exposed TG-rich ssDNA, telomere clustering in Cajal bodies, and telomere bouquet development during spindle formation, where proximity could raise the local concentration of telomeric ssDNA favouring Cdc13 DDE toward the longest ssDNAs available (95–97).

## DATA AVAILABILITY

The data underlying this article are available in the article, in its online supplementary material, or will be shared upon reasonable request to the corresponding author.

## SUPPLEMENTARY DATA

Supplementary Data are available at NAR online.

## AUTHOR CONTRIBUTIONS

David Nickens: Conceptualization, Formal analysis, Methodology, Validation, Protein purification (Cdc13 and Pif1), Writing—original draft. Zhitong Feng: Protein purification (OB1, OB3, and Cdc13ΔOB1), Writing—review & editing. Jiangchuan Shen: Protein purification (RPA), Writing—review & editing. Spencer J. Gray: Methodology, Writing—review & editing. Robert Simmons: Protein purification (Hrq1), Writing—review & editing. Hengyao Niu: Review & editing. Matthew Bochman: Conceptualization, Formal analysis, Methodology, Validation, Writing— review & editing.

## ACKNOWLEDGEMENTS

We thank Deborah Wuttke for the Cdc13 DBD expression plasmid, as well as Quan Wang for the pFastBac-Cdc13 plasmid and for his helpful advice with insect cell culture.

## REFERENCES

1. Herrmann, M., Pusceddu, I., Marz, W. and Herrmann, W. (2018) Telomere biology and age-related diseases. Clin Chem Lab Med, 56, 1210–1222.

2. Wellinger, R.J. and Zakian, V.A. (2012) Everything you ever wanted to know about Saccharomyces cerevisiae telomeres: beginning to end. Genetics, 191, 1073–1105.

3. Gao, H., Cervantes, R.B., Mandell, E.K., Otero, J.H. and Lundblad, V. (2007) RPA-like proteins mediate yeast telomere function. Nat Struct Mol Biol.

4. Bochman, M.L., Paeschke, K., Chan, A. and Zakian, V.A. (2014) Hrq1, a homolog of the human RecQ4 helicase, acts catalytically and structurally to promote genome integrity. Cell Rep, 6, 346–356.

5. Nickens, D.G. and Bochman, M.L. (2022) Genetic and biochemical interactions of yeast DNA helicases. Methods, 204, 234–240.

6. Nickens, D.G., Rogers, C.M. and Bochman, M.L. (2018) The Saccharomyces cerevisiae Hrq1 and Pif1 DNA helicases synergistically modulate telomerase activity in vitro. J Biol Chem, 293, 14481–14496.

7. Bochman, M.L., Sabouri, N. and Zakian, V.A. (2010) Unwinding the functions of the Pif1 family helicases. DNA Repair (Amst), 9, 237–249.

8. Caldwell, C.C. and Spies, M. (2020) Dynamic elements of replication protein A at the crossroads of DNA replication, recombination, and repair. Crit Rev Biochem Mol Biol, 55, 482–507.

9. de Laat, W.L., Appeldoorn, E., Sugasawa, K., Weterings, E., Jaspers, N.G. and Hoeijmakers, J.H. (1998) DNA-binding polarity of human replication protein A positions nucleases in nucleotide excision repair. Genes Dev, 12, 2598–2609.

10. Iftode, C. and Borowiec, J.A. (2000) 5’ --> 3’ molecular polarity of human replication protein A (hRPA) binding to pseudo-origin DNA substrates. Biochemistry, 39, 11970–11981.

11. Fanning, E., Klimovich, V. and Nager, A.R. (2006) A dynamic model for replication protein A (RPA) function in DNA processing pathways. Nucleic Acids Res, 34, 4126–4137.

12. Gan, X., Zhang, Y., Jiang, D., Shi, J., Zhao, H., Xie, C., Wang, Y., Xu, J., Zhang, X., Cai, G. et al. (2023) Proper RPA acetylation promotes accurate DNA replication and repair. Nucleic Acids Res, 51, 5565–5583.

13. Mondal, A. and Bhattacherjee, A. (2020) Mechanism of Dynamic Binding of Replication Protein A to ssDNA. J Chem Inf Model, 60, 5057–5069.

14. Nguyen, B., Sokoloski, J., Galletto, R., Elson, E.L., Wold, M.S. and Lohman, T.M. (2014) Diffusion of human replication protein A along single-stranded DNA. J Mol Biol, 426, 3246–3261.

15. Pokhrel, N., Caldwell, C.C., Corless, E.I., Tillison, E.A., Tibbs, J., Jocic, N., Tabei, S.M.A., Wold, M.S., Spies, M. and Antony, E. (2019) Dynamics and selective remodeling of the DNA-binding domains of RPA. Nat Struct Mol Biol, 26, 129–136.

16. Mersch, K.N., Sokoloski, J.E., Nguyen, B., Galletto, R. and Lohman, T.M. (2023) “Helicase” Activity promoted through dynamic interactions between a ssDNA translocase and a diffusing SSB protein. Proc Natl Acad Sci U S A, 120, e2216777120.

17. Chen, R., Subramanyam, S., Elcock, A.H., Spies, M. and Wold, M.S. (2016) Dynamic binding of replication protein a is required for DNA repair. Nucleic Acids Res, 44, 5758–5772.

18. Alani, E., Thresher, R., Griffith, J.D. and Kolodner, R.D. (1992) Characterization of DNA-binding and strand-exchange stimulation properties of y-RPA, a yeast single-strand-DNA-binding protein. J Mol Biol, 227, 54–71.

19. Schramke, V., Luciano, P., Brevet, V., Guillot, S., Corda, Y., Longhese, M.P., Gilson, E. and Geli, V. (2004) RPA regulates telomerase action by providing Est1p access to chromosome ends. Nat Genet, 36, 46–54.

20. Luciano, P., Coulon, S., Faure, V., Corda, Y., Bos, J., Brill, S.J., Gilson, E., Simon, M.N. and Geli, V. (2012) RPA facilitates telomerase activity at chromosome ends in budding and fission yeasts. Embo J, 31, 2034–2046.

21. Greetham, M., Skordalakes, E., Lydall, D. and Connolly, B.A. (2015) The Telomere Binding Protein Cdc13 and the Single-Stranded DNA Binding Protein RPA Protect Telomeric DNA from Resection by Exonucleases. J Mol Biol, 427, 3023–3030.

22. Audry, J., Maestroni, L., Delagoutte, E., Gauthier, T., Nakamura, T.M., Gachet, Y., Saintome, C., Geli, V. and Coulon, S. (2015) RPA prevents G-rich structure formation at lagging-strand telomeres to allow maintenance of chromosome ends. Embo J, 34, 1942–1958.

23. Maestroni, L., Audry, J., Luciano, P., Coulon, S., Geli, V. and Corda, Y. (2020) RPA and Pif1 cooperate to remove G-rich structures at both leading and lagging strand. Cell Stress, 4, 48–63.

24. Nugent, C.I., Hughes, T.R., Lue, N.F. and Lundblad, V. (1996) Cdc13p: a single-strand telomeric DNA-binding protein with a dual role in yeast telomere maintenance. Science, 274, 249–252.

25. Qi, H. and Zakian, V.A. (2000) The Saccharomyces telomere-binding protein Cdc13p interacts with both the catalytic subunit of DNA polymerase alpha and the telomerase-associated est1 protein. Genes Dev, 14, 1777–1788.

26. Chandra, A., Hughes, T.R., Nugent, C.I. and Lundblad, V. (2001) Cdc13 both positively and negatively regulates telomere replication. Genes Dev, 15, 404–414.

27. Pennock, E., Buckley, K. and Lundblad, V. (2001) Cdc13 delivers separate complexes to the telomere for end protection and replication. Cell, 104, 387–396.

28. Zappulla, D.C., Roberts, J.N., Goodrich, K.J., Cech, T.R. and Wuttke, D.S. (2009) Inhibition of yeast telomerase action by the telomeric ssDNA-binding protein, Cdc13p. Nucleic Acids Res, 37, 354–367.

29. Giraud-Panis, M.J., Teixeira, M.T., Geli, V. and Gilson, E. (2010) CST meets shelterin to keep telomeres in check. Mol Cell, 39, 665–676.

30. Ge, Y., Wu, Z., Chen, H., Zhong, Q., Shi, S., Li, G., Wu, J. and Lei, M. (2020) Structural insights into telomere protection and homeostasis regulation by yeast CST complex. Nat Struct Mol Biol, 27, 752–762.

31. Wu, Y. and Zakian, V.A. (2011) The telomeric Cdc13 protein interacts directly with the telomerase subunit Est1 to bring it to telomeric DNA ends in vitro. Proc Natl Acad Sci U S A, 108, 20362–20369.

32. Liu, C.C., Gopalakrishnan, V., Poon, L.F., Yan, T. and Li, S. (2014) Cdk1 regulates the temporal recruitment of telomerase and Cdc13-Stn1-Ten1 complex for telomere replication. Mol Cell Biol, 34, 57–70.

33. Calvo, O., Grandin, N., Jordan-Pla, A., Minambres, E., Gonzalez-Polo, N., Perez-Ortin, J.E. and Charbonneau, M. (2019) The telomeric Cdc13-Stn1-Ten1 complex regulates RNA polymerase II transcription. Nucleic Acids Res, 47, 6250–6268.

34. Langston, R.E., Palazzola, D., Bonnell, E., Wellinger, R.J. and Weinert, T. (2020) Loss of Cdc13 causes genome instability by a deficiency in replication-dependent telomere capping. PLoS Genet, 16, e1008733.

35. Lin, Y.Y., Li, M.H., Chang, Y.C., Fu, P.Y., Ohniwa, R.L., Li, H.W. and Lin, J.J. (2021) Dynamic DNA Shortening by Telomere-Binding Protein Cdc13. J Am Chem Soc, 143, 5815–5825.

36. Tseng, S.F., Lin, J.J. and Teng, S.C. (2006) The telomerase-recruitment domain of the telomere binding protein Cdc13 is regulated by Mec1p/Tel1p-dependent phosphorylation. Nucleic Acids Res, 34, 6327–6336.

37. Vodenicharov, M.D. and Wellinger, R.J. (2006) DNA degradation at unprotected telomeres in yeast is regulated by the CDK1 (Cdc28/Clb) cell-cycle kinase. Mol Cell, 24, 127–137.

38. Wu, Y., DiMaggio, P.A., Jr., Perlman, D.H., Zakian, V.A. and Garcia, B.A. (2013) Novel phosphorylation sites in the S. cerevisiae Cdc13 protein reveal new targets for telomere length regulation. J Proteome Res, 12, 316–327.

39. Gopalakrishnan, V., Tan, C.R. and Li, S. (2017) Sequential phosphorylation of CST subunits by different cyclin-Cdk1 complexes orchestrate telomere replication. Cell Cycle, 16, 1271–1287.

40. Bianchi, A., Negrini, S. and Shore, D. (2004) Delivery of yeast telomerase to a DNA break depends on the recruitment functions of Cdc13 and Est1. Mol Cell, 16, 139–146.

41. Chen, Y.F., Lu, C.Y., Lin, Y.C., Yu, T.Y., Chang, C.P., Li, J.R., Li, H.W. and Lin, J.J. (2016) Modulation of yeast telomerase activity by Cdc13 and Est1 in vitro. Sci Rep, 6, 34104.

42. Hang, L.E., Liu, X., Cheung, I., Yang, Y. and Zhao, X. (2011) SUMOylation regulates telomere length homeostasis by targeting Cdc13. Nat Struct Mol Biol, 18, 920–926.

43. Sun, J., Yang, Y., Wan, K., Mao, N., Yu, T.Y., Lin, Y.C., DeZwaan, D.C., Freeman, B.C., Lin, J.J., Lue, N.F. et al. (2011) Structural bases of dimerization of yeast telomere protein Cdc13 and its interaction with the catalytic subunit of DNA polymerase alpha. Cell Res, 21, 258–274.

44. Olson, C.L., Barbour, A.T. and Wuttke, D.S. (2022) Filling in the blanks: how the C-strand catches up to the G-strand at replicating telomeres using CST. Nat Struct Mol Biol, 29, 730–733.

45. Mitton-Fry, R.M., Anderson, E.M., Theobald, D.L., Glustrom, L.W. and Wuttke, D.S. (2004) Structural basis for telomeric single-stranded DNA recognition by yeast Cdc13. J Mol Biol, 338, 241–255.

46. Theobald, D.L., Mitton-Fry, R.M. and Wuttke, D.S. (2003) Nucleic acid recognition by OB-fold proteins. Annu Rev Biophys Biomol Struct, 32, 115–133.

47. Lewis, K.A., Pfaff, D.A., Earley, J.N., Altschuler, S.E. and Wuttke, D.S. (2014) The tenacious recognition of yeast telomere sequence by Cdc13 is fully exerted by a single OB-fold domain. Nucleic Acids Res, 42, 475–484.

48. Mersaoui, S.Y., Bonnell, E. and Wellinger, R.J. (2018) Nuclear import of Cdc13 limits chromosomal capping. Nucleic Acids Res, 46, 2975–2989.

49. Lin, J.J. and Zakian, V.A. (1996) The Saccharomyces CDC13 protein is a single-strand TG1-3 telomeric DNA-binding protein in vitro that affects telomere behavior in vivo. Proc Natl Acad Sci U S A, 93, 13760–13765.

50. Lin, Y.C., Hsu, C.L., Shih, J.W. and Lin, J.J. (2001) Specific binding of single-stranded telomeric DNA by Cdc13p of Saccharomyces cerevisiae. J Biol Chem, 276, 24588–24593.

51. Glustrom, L.W., Lyon, K.R., Paschini, M., Reyes, C.M., Parsonnet, N.V., Toro, T.B., Lundblad, V. and Wuttke, D.S. (2018) Single-stranded telomere-binding protein employs a dual rheostat for binding affinity and specificity that drives function. Proc Natl Acad Sci U S A, 115, 10315–10320.

52. Anderson, E.M., Halsey, W.A. and Wuttke, D.S. (2002) Delineation of the high-affinity single-stranded telomeric DNA-binding domain of Saccharomyces cerevisiae Cdc13. Nucleic Acids Res, 30, 4305–4313.

53. Mitchell, M.T., Smith, J.S., Mason, M., Harper, S., Speicher, D.W., Johnson, F.B. and Skordalakes, E. (2010) Cdc13 N-terminal dimerization, DNA binding, and telomere length regulation. Mol Cell Biol, 30, 5325–5334.

54. Hughes, T.R., Weilbaecher, R.G., Walterscheid, M. and Lundblad, V. (2000) Identification of the single-strand telomeric DNA binding domain of the Saccharomyces cerevisiae Cdc13 protein. Proc Natl Acad Sci U S A, 97, 6457–6462.

55. Eldridge, A.M., Halsey, W.A. and Wuttke, D.S. (2006) Identification of the determinants for the specific recognition of single-strand telomeric DNA by Cdc13. Biochemistry, 45, 871–879.

56. Eldridge, A.M. and Wuttke, D.S. (2008) Probing the mechanism of recognition of ssDNA by the Cdc13-DBD. Nucleic Acids Res, 36, 1624–1633.

57. Mandell, E.K., Gelinas, A.D., Wuttke, D.S. and Lundblad, V. (2011) Sequence-specific binding to telomeric DNA is not a conserved property of the Cdc13 DNA binding domain. Biochemistry, 50, 6289–6291.

58. Mason, M., Wanat, J.J., Harper, S., Schultz, D.C., Speicher, D.W., Johnson, F.B. and Skordalakes, E. (2013) Cdc13 OB2 dimerization required for productive Stn1 binding and efficient telomere maintenance. Structure, 21, 109–120.

59. Azvolinsky, A., Dunaway, S., Torres, J., Bessler, J. and Zakian, V.A. (2006) The S. cerevisiae Rrm3p DNA helicase moves with the replication fork and affects replication of all yeast chromosomes. Genes Dev, 20, 3104–3116.

60. Azvolinsky, A., Dunaway, S., Torres, J.Z., Bessler, J.B. and Zakian, V.A. (2006) The S. cerevisiae Rrm3p DNA helicase moves with the replication fork and affects replication of all yeast chromosomes. Genes Dev, 20, 3104–3116.

61. Tran, P.L.T., Pohl, T.J., Chen, C.F., Chan, A., Pott, S. and Zakian, V.A. (2017) PIF1 family DNA helicases suppress R-loop mediated genome instability at tRNA genes. Nat Commun, 8, 15025.

62. Pennaneach, V., Putnam, C.D. and Kolodner, R.D. (2006) Chromosome healing by de novo telomere addition in Saccharomyces cerevisiae. Mol Microbiol, 59, 1357–1368.

63. Dewar, J.M. and Lydall, D. (2012) Similarities and differences between “uncapped” telomeres and DNA double-strand breaks. Chromosoma, 121, 117–130.

64. Strecker, J., Stinus, S., Caballero, M.P., Szilard, R.K., Chang, M. and Durocher, D. (2017) A sharp Pif1-dependent threshold separates DNA double-strand breaks from critically short telomeres. Elife, 6.

65. Chib, S., Griffin, W.C., Gao, J., Proffitt, D.R., Byrd, A.K. and Raney, K.D. (2023) Pif1 Helicase Mediates Remodeling of Protein-Nucleic Acid Complexes by Promoting Dissociation of Sub1 from G-Quadruplex DNA and Cdc13 from G-Rich Single-Stranded DNA. Biochemistry.

66. Nickens, D.G., Sausen, C.W. and Bochman, M.L. (2019) The Biochemical Activities of the Saccharomyces cerevisiae Pif1 Helicase Are Regulated by Its N-Terminal Domain. Genes (Basel), 10.

67. Phillips, J.A., Chan, A., Paeschke, K. and Zakian, V.A. (2015) The pif1 helicase, a negative regulator of telomerase, acts preferentially at long telomeres. PLoS Genet, 11, e1005186.

68. Stinus, S., Paeschke, K. and Chang, M. (2018) Telomerase regulation by the Pif1 helicase: a length-dependent effect? Curr Genet, 64, 509–513.

69. Shen, J., Zhao, Y., Pham, N.T., Li, Y., Zhang, Y., Trinidad, J., Ira, G., Qi, Z. and Niu, H. (2022) Deciphering the mechanism of processive ssDNA digestion by the Dna2-RPA ensemble. Nat Commun, 13, 359.

70. Rogers, C.M., Lee, C.Y., Parkins, S., Buehler, N.J., Wenzel, S., Martinez-Marquez, F., Takagi, Y., Myong, S. and Bochman, M.L. (2020) The yeast Hrq1 helicase stimulates Pso2 translesion nuclease activity and thereby promotes DNA interstrand crosslink repair. J Biol Chem, 295, 8945–8957.

71. Wieser, T.A. and Wuttke, D.S. (2022) Replication Protein A Utilizes Differential Engagement of Its DNA-Binding Domains to Bind Biologically Relevant ssDNAs in Diverse Binding Modes. Biochemistry, 61, 2592–2606.

72. Bochman, M.L. and Schwacha, A. (2008) The Mcm2-7 complex has in vitro helicase activity. Mol Cell, 31, 287–293.

73. Barrows, J.K. and Van Dyke, M.W. (2022) Biolayer interferometry for DNA-protein interactions. PLoS One, 17, e0263322.

74. Lewis, K.A. and Wuttke, D.S. (2012) Telomerase and telomere-associated proteins: structural insights into mechanism and evolution. Structure, 20, 28–39.

75. Kozlov, A.G. and Lohman, T.M. (2002) Kinetic mechanism of direct transfer of Escherichia coli SSB tetramers between single-stranded DNA molecules. Biochemistry, 41, 11611–11627.

76. Jurikova, K., Gajarsky, M., Hajikazemi, M., Nosek, J., Prochazkova, K., Paeschke, K., Trantirek, L. and Tomaska, L. (2020) Role of folding kinetics of secondary structures in telomeric G-overhangs in the regulation of telomere maintenance in Saccharomyces cerevisiae. J Biol Chem, 295, 8958–8971.

77. Tran, P.L., Mergny, J.L. and Alberti, P. (2011) Stability of telomeric G-quadruplexes. Nucleic Acids Res, 39, 3282–3294.

78. Dickey, T.H., Altschuler, S.E. and Wuttke, D.S. (2013) Single-stranded DNA-binding proteins: multiple domains for multiple functions. Structure, 21, 1074–1084.

79. Rice, C. and Skordalakes, E. (2016) Structure and function of the telomeric CST complex. Comput Struct Biotechnol J, 14, 161–167.

80. Itriago, H., Jaiswal, R.K., Philipp, S. and Cohn, M. (2022) The telomeric 5’ end nucleotide is regulated in the budding yeast Naumovozyma castellii. Nucleic Acids Res, 50, 281–292.

81. Hoerr, R.E., Eng, A., Payen, C., Di Rienzi, S.C., Raghuraman, M.K., Dunham, M.J., Brewer, B.J. and Friedman, K.L. (2023) Hotspot of de novo telomere addition stabilizes linear amplicons in yeast grown in sulfate-limiting conditions. Genetics.

82. Anand, R.P., Shah, K.A., Niu, H., Sung, P., Mirkin, S.M. and Freudenreich, C.H. (2012) Overcoming natural replication barriers: differential helicase requirements. Nucleic Acids Res, 40, 1091–1105.

83. Byrd, A.K. and Raney, K.D. (2017) Structure and function of Pif1 helicase. Biochem Soc Trans, 45, 1159–1171.

84. Rogers, C.M. and Bochman, M.L. (2017) Saccharomyces cerevisiae Hrq1 helicase activity is affected by the sequence but not the length of single-stranded DNA. Biochem Biophys Res Commun, 486, 1116–1121.

85. Rogers, C.M., Wang, J.C., Noguchi, H., Imasaki, T., Takagi, Y. and Bochman, M.L. (2017) Yeast Hrq1 shares structural and functional homology with the disease-linked human RecQ4 helicase. Nucleic Acids Res, 45, 5217–5230.

86. Antony, E. and Lohman, T.M. (2019) Dynamics of E. coli single stranded DNA binding (SSB) protein-DNA complexes. Semin Cell Dev Biol, 86, 102–111.

87. Qi, H. and Zakian, V.A. (2000) The Saccharomyces telomere-binding protein Cdc13p interacts with both the catalytic subunit of DNA polymerase α and the telomerase-associated Est1 protein. Genes Dev., 14, 1777–1788.

88. Chandra, A., Hughes, T.R., Nugent, C.I. and Lundblad, V. (2001) Cdc13 both positively and negatively regulates telomere replication. Genes Dev., 15, 404–414.

89. Lue, N.F., Chan, J., Wright, W.E. and Hurwitz, J. (2014) The CDC13-STN1-TEN1 complex stimulates Pol alpha activity by promoting RNA priming and primase-to-polymerase switch. Nat Commun, 5, 5762.

90. Tseng, S., Lin, J. and Teng, S. (2006) The telomerase-recruitment domain of the telomere binding protein Cdc13 is regulated by Mec1p/Tel1p-dependent phosphorylation. Nucl Acids Res, 34, 6327–6336.

91. Vodenicharov, M. and Wellinger, R.J. (2006) DNA degradation at unprotected telomeres in yeast is regulated by the CDK1 (Cdc28/Clb) cell-cycle kinase. Mol Cell, 24, 127–137.

92. Ononye, O.E., Sausen, C.W., Balakrishnan, L. and Bochman, M.L. (2020) Lysine Acetylation Regulates the Activity of Nuclear Pif1. J Biol Chem.

93. Ononye, O.E., Sausen, C.W., Bochman, M.L. and Balakrishnan, L. (2020) Dynamic regulation of Pif1 acetylation is crucial to the maintenance of genome stability. Curr Genet.

94. Hemphill, W.O., Voong, C.K., Fenske, R., Goodrich, J.A. and Cech, T.R. (2023) Multiple RNA- and DNA-binding proteins exhibit direct transfer of polynucleotides with implications for target-site search. Proc Natl Acad Sci U S A, 120, e2220537120.

95. Bianchi, A., Negrini, S. and Shore, D. (2004) Delivery of yeast telomerase to a DNA break depends on the recruitment functions of Cdc13 and Est1. Mol. Cell, 16, 139–146.

96. Moiseeva, V., Amelina, H., Collopy, L.C., Armstrong, C.A., Pearson, S.R. and Tomita, K. (2017) The telomere bouquet facilitates meiotic prophase progression and exit in fission yeast. Cell Discov, 3, 17041.

97. Zhong, F.L., Batista, L.F., Freund, A., Pech, M.F., Venteicher, A.S. and Artandi, S.E. (2012) TPP1 OB-fold domain controls telomere maintenance by recruiting telomerase to chromosome ends. Cell, 150, 481–494.

