## Supplemental material for "Cdc13 exhibits dynamic DNA strand exchange in the presence of telomeric DNA"

**Supplementary Materials**

**Supplementary tables**

**Table S1. Oligonucleotides used in this study.**

| **Number** | **Name** | **Sequence** |
| --- | --- | --- |
| MB915 | Tel30G | GTGTGGGTGTGGTGTGGGTGTGGTGTGGGT |
| MB464 | Poly(dT)_30_ | TTTTTTTTTTTTTTTTTTTTTTTTTTTTTT |
| MB1169 | Tel25G fork | GGTGTGGTGTGGGTGTGGTGTGGGTGTGTCACTCACATAGCGTTC |
| MB1170 | Tel25G fork | GAACGCTATGTGAGTGACACTGGGTGTGGTGTGGGTGTGGTGTGG |
| MB1772 | Poly(T)25 fork | /5IRD700/GAACGCTATGTGAGTGACACTTTTTTTTTTTTTTTTTTTTTTTTT |
| MB1013 | Tel15G | TGTGGTGTGTGTGGG |
| MB1117 | Tel50G | GTGTGGGTGTGGTGTGGGTGTGGTGTGGGTGTGTGGGTGTGGTGTGGGTG |
| MB1755 | IR700-Poly(dT)_30_ | /5IRD700/TTTTTTTTTTTTTTTTTTTTTTTTTTTTTT |
| MB1929 | IR700-Tel15G | /5IRD700/TGTGGTGTGTGTGGG |
| MB1930 | IR700-Tel30G | /5IRD700/CGCCATGCTGATCCGTGTGGTGTGTGTGGG |
| MB1931 | IR700-Tel50G | /5IRD700/GTGTGGGTGTGGTGTGGGTGTGGTGTGGGTGTGTGGTGTGGTGTGTGTG |
| MB1981 | IR800-Tel30G | /5IRD800/CGCCATGCTGATCCGTGTGGTGTGTGTGGG |
| MB1982 | IR800-Poly(dT)_30_ | /5IRD800/TTTTTTTTTTTTTTTTTTTTTTTTTTTTTT |
| MB1983 | Tel11G | GTGTGGGTGTG |
| MB2018 | IR700-Tel11G | /5IRD700/GTGTGGGTGTG |
| MB2081 | Biotinylated Tel30G | /5BiotiTEG/ CGCCATGCTGATCCGTGTGGTGTGTGTGGG |

**Supplementary figure legends**

**Figure S1. Purified protein preparations.** A) Insect cells were used to generate full-length wild-type Cdc13, which was used in all assays where noted, and the Hrq1 helicase. B) *Escherichia coli* was also used to make full-length wild-type Cdc13, which was only used in Figure 3B, and the Pif1 helicase. C) *E. coli* was likewise used to generate yeast RPA (yRPA) and the Cdc13 truncations: OB1, OB3, and Cdc13ΔOB1 (OB1Δ). All protein preparations were separated on SDS-PAGE gels and stained with Coomassie.

**Figure S2. AlphaFold predictions of Cdc13 structure.** A) AlphaFold predicts that the tertiary structure of Cdc13 is dominated by three globular domains comprised of OB1, OB3, and OB2/4 connected by disordered linkers. B) AlphaFold Multimer model of a Cdc13 dimer. One subunit is shown in purple, and the other is pink. C) The same model as in (B) but coloured by domain. The OB1 domains are red and dark red, the OB2s are yellow and dark orange, the OB3s are lime and green, and the OB4s are cyan and blue in monomers 1 and 2, respectively. D) Predicted alignment error plot generated for the Cdc13 dimer by AlphaFold Multimer. The folding of the OB motifs and their interactions were predicted with high confidence. AlphaFold Multimer predicts that OB2 and OB4 interact with each other in each monomer, as well as between subunits in the dimer. The OB1 domains are also predicted to interact between the monomers in the dimer, but the OB3 domains are devoid of predicted interactions both within and between monomers.

**Figure S3. Model for the disruption of Cdc13-ssDNA complexes by DNA helicases.** A) The assay is based on a Cdc13-ssDNA gel shift. If a helicase can disrupt the Cdc13-ssDNA complex, then titrating in the helicase should decrease the amount of Cdc13-bound ssDNA in a helicase concentration-dependent manner. B) Because Cdc13, Hrq1, and Pif1 are all ssDNA binding proteins, a protein trap was added to the assay diagrammed in (A) to prevent rebinding of the proteins to the substrate. Here, the protein trap is a molar excess of either a differentially labelled ssDNA or an unlabelled ssDNA.

**Figure S4. The isolated OB3 domain does not demonstrate DDE.** Representative DDE assay for OB3. Recombinant OB3 (7.5 nM monomer) was prebound to 2 nM IR700-Tel11G and then exposed to increasing concentrations (0.4-50 nM) of unlabelled Tel11G competitor substrate. No DDE or competition was observed. The lane marked ‘B’ contains OB3 and IR700-Tel11G but was boiled prior to loading on the gel, and thus it serves as a control for where unbound Tel11G would migrate through the gel.

**Figure S5. The isolated OB1 domain has low ssDNA binding affinity and does not demonstrate DDE.** A) Representative gel shift assay using labelled Tel30G ssDNA and increasing concentrations (7.8-1000 nM) of recombinant OB1. The *k_1/2_* calculated from triplicate assays (830 nM) indicates that OB1 has low DNA binding affinity compared to full-length Cdc13 (*K_D_* = 3.7 nM). B) Representative DDE assay for OB1. A high concentration of labelled Tel30G (1 μM) was used to achieve 100% binding by OB1 in the absence of competitor, whereas the typical DDE assay with full-length Cdc13 requires only 2 nM ssDNA substrate. A 0.16-2.5 μM range of competitor DNA concentrations was tested.

**Figure S6. Example BLI assays to observe DDE.** In a typical assay, biotinylated Tel30G ssDNA was immobilized on a streptavidin-coated BLI sensor, washed, exposed to Cdc13 for ssDNA binding (Association), exposed to a large volume of buffer to allow dissociation of Cdc13 from the ssDNA, and then exposed to a large volume of buffer containing a molar excess of competitor ssDNA to allow DDE to occur.

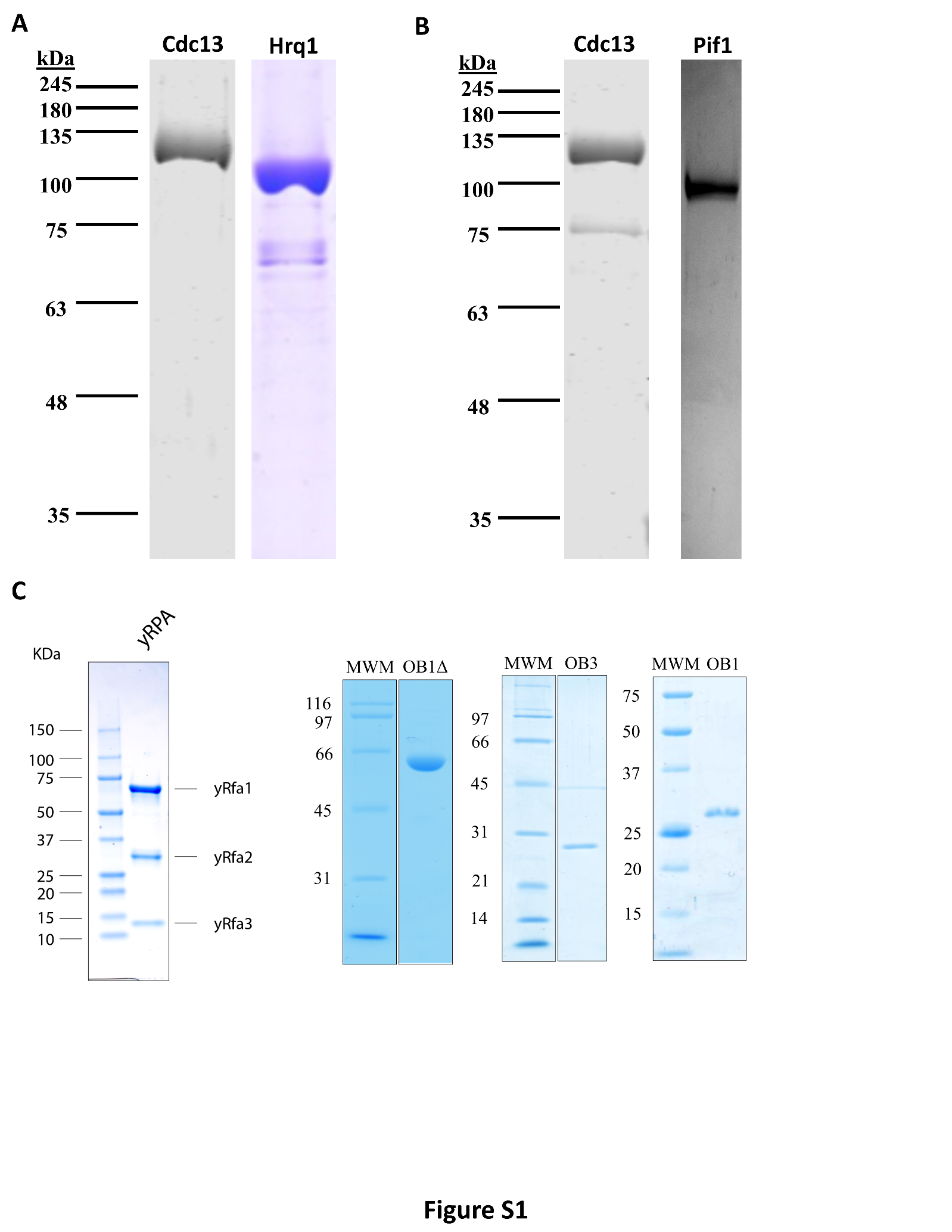

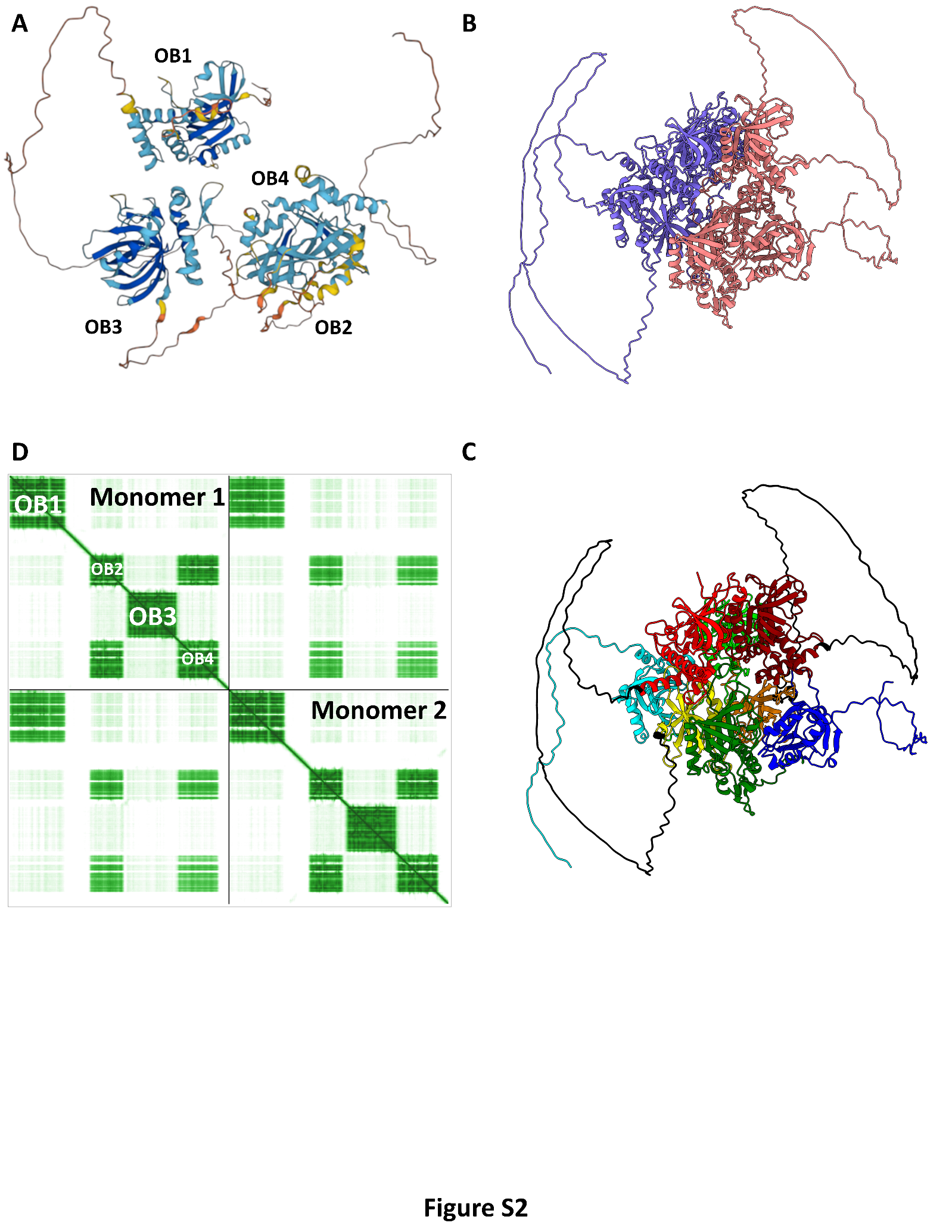

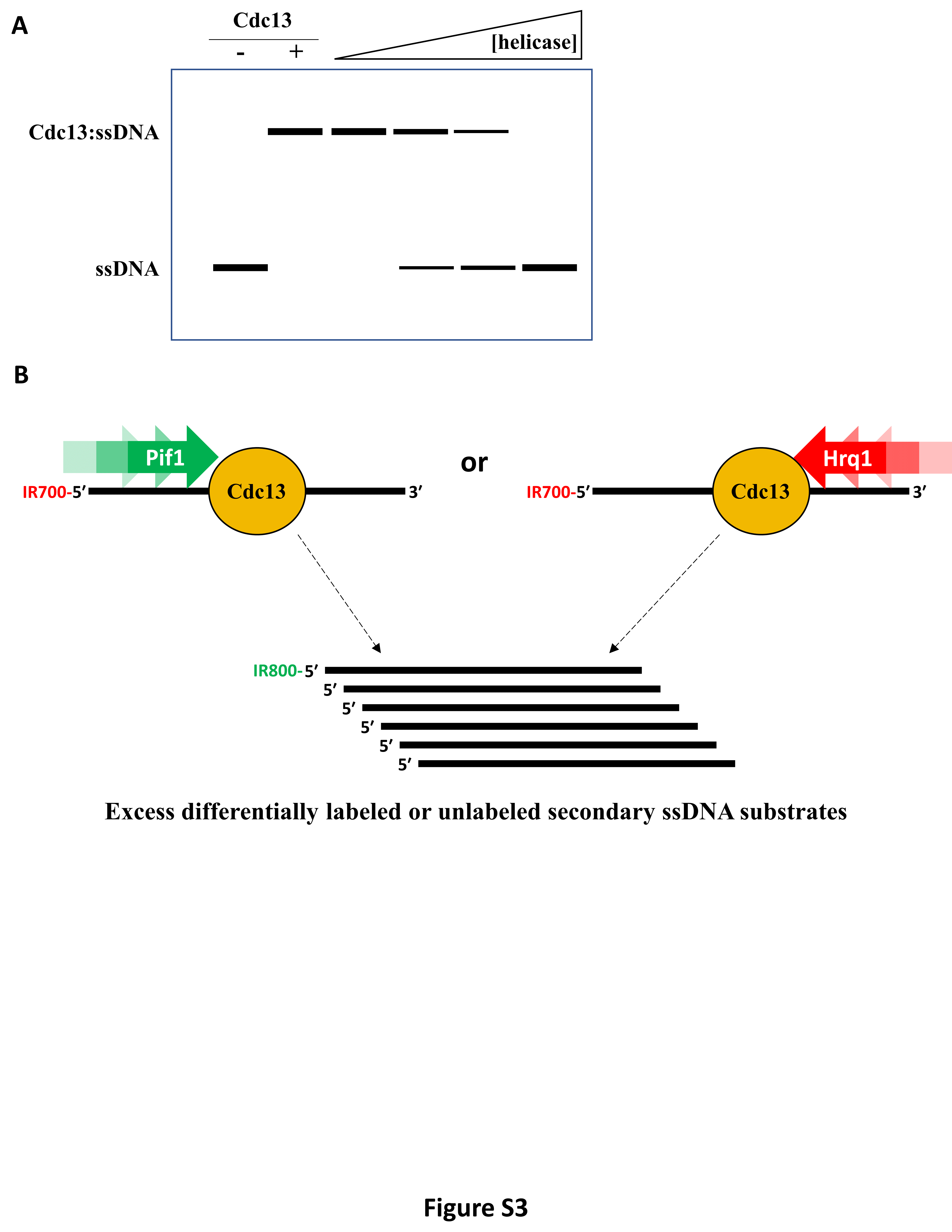

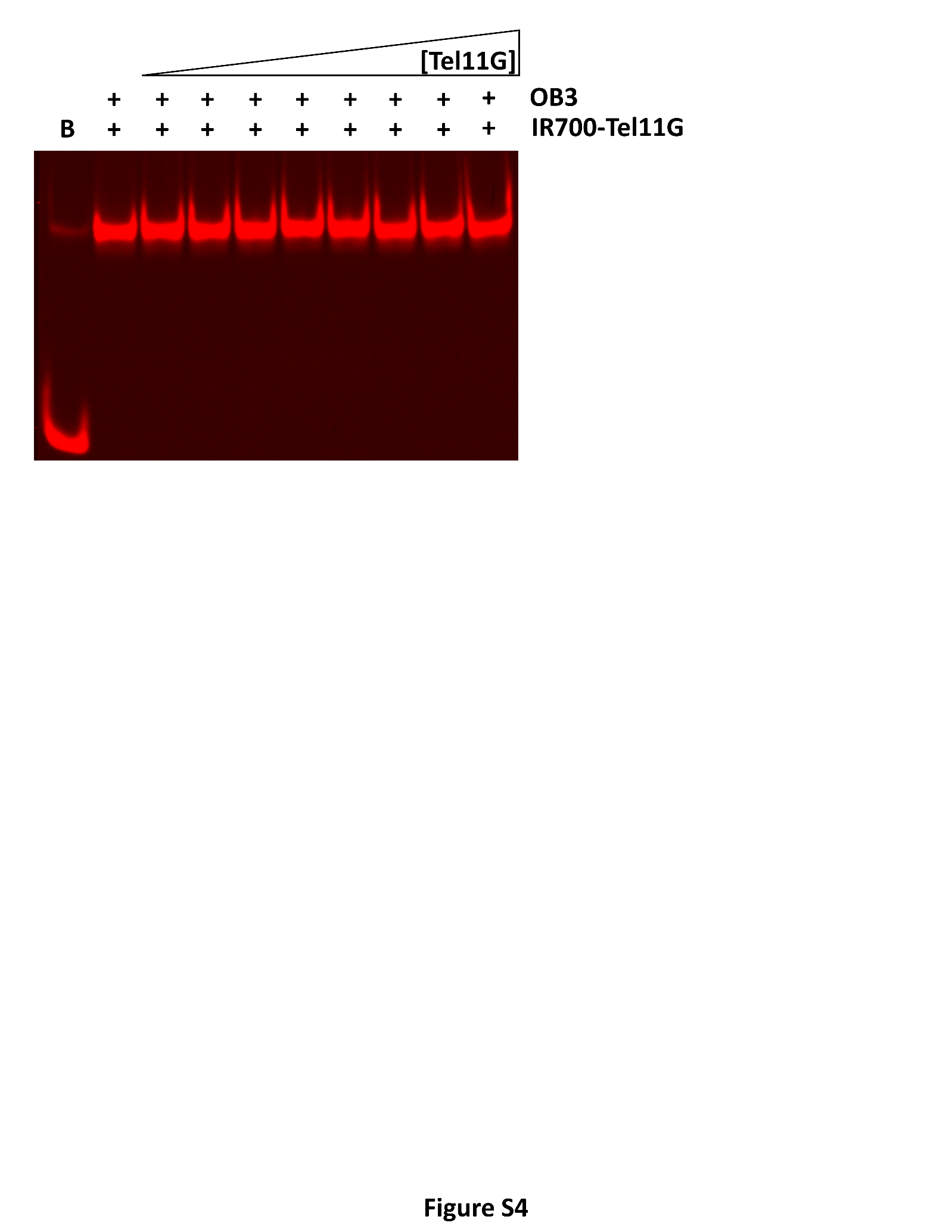

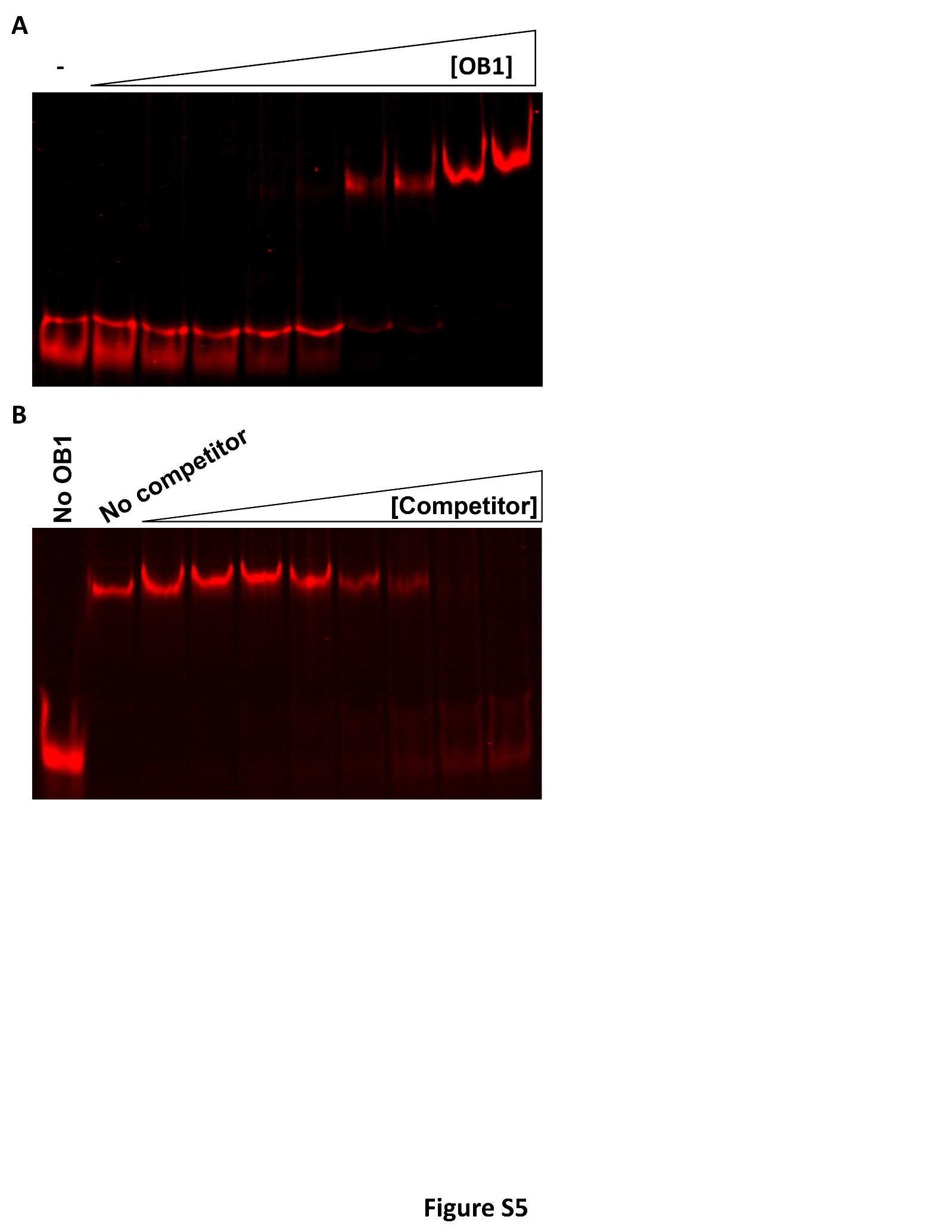

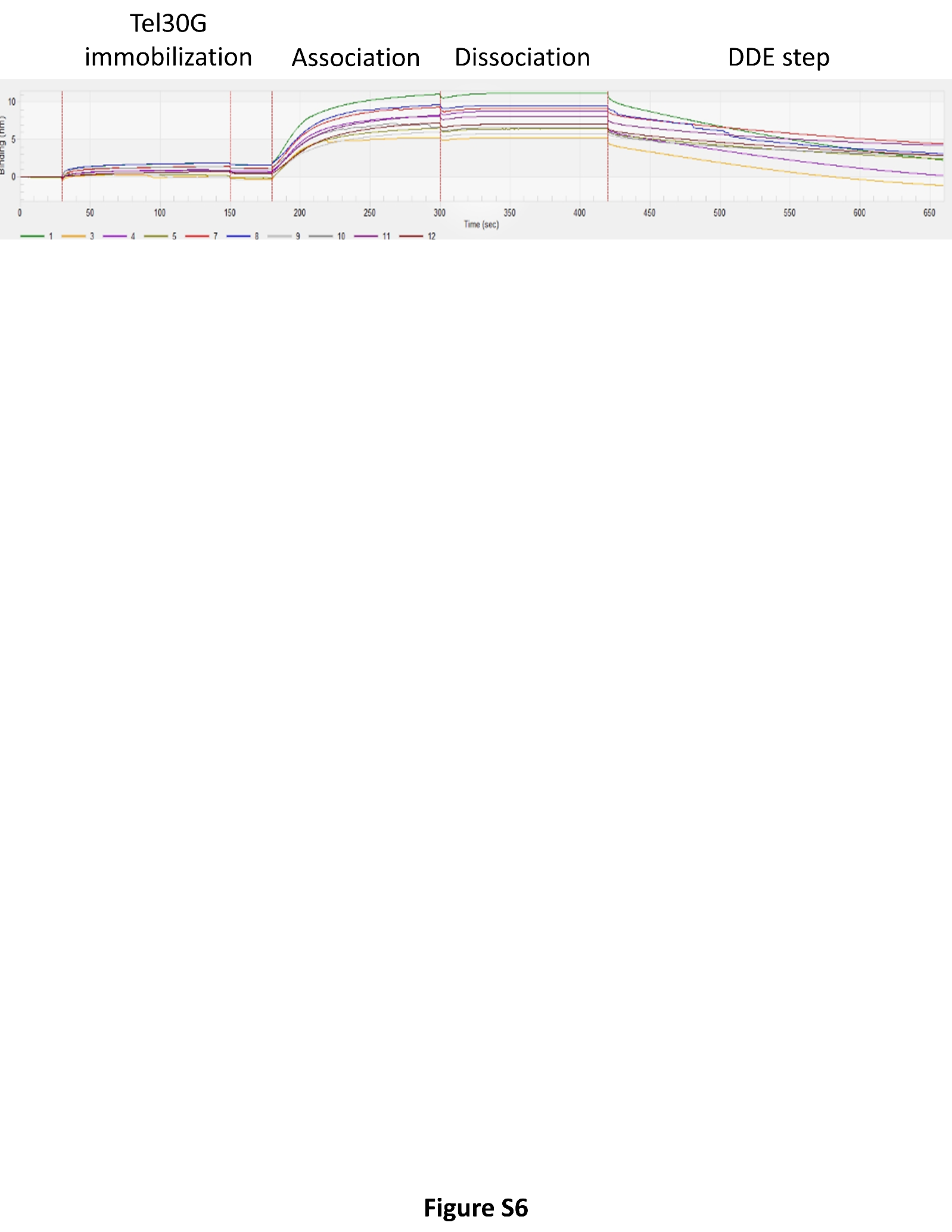
